# Frequency modulated timer regulates mammalian hibernation

**DOI:** 10.1101/2021.11.12.468369

**Authors:** Shingo Gibo, Yoshifumi Yamaguchi, Gen Kurosawa

## Abstract

Mammalian hibernators decrease basal metabolism and body temperature (Tb) to minimize energy expenditure in harsh seasons. During hibernation, Tb drops to low temperature (<10 °C) and remains constant for days, known as deep torpor in small mammalian hibernators. Spontaneous interbout arousals interrupt torpor bouts, when Tb recovers to euthermic state ~37 °C. Torpor-interbout arousal event repeats during several months of hibernation. However, little is known about mechanisms governing Tb fluctuation across torpor-interbout arousal cycles during hibernation. Recent improvement in data-logging techniques enables us to monitor Tb for more than hundred days with high precision, opening up new avenues for quantitative analysis to address the principle governing Tb fluctuation. Here, we analyzed Tb fluctuation across torpor-interbout arousal cycle of Syrian hamster, which can hibernate in response to chronic cold and short photoperiod under a laboratory condition, using generalized harmonic analysis and discovered a model with frequency modulation quantitatively reproducing Tb fluctuation. This analysis also identified an unexpectedly longer period of 120–430 days as the period that modulates another period of several days, generating Tb fluctuation for Syrian hamster. We propose that concerted action of two endogenous periods governs torpor-interbout arousal cycles during hibernation.

## Introduction

Hibernation is a strategy for organisms to survive in the environment with limited food and water availability (Geiser, 2013) (Mohr et al., 2020). During a season with little or no food, small mammalian hibernators drastically decrease their basal metabolism and core body temperature (Tb) to 10 °C and become immobile. This hypometabolic, hypothermic, and immobile state is called deep torpor. Interbout arousal (IBA) interrupt deep torpor, during which Tb rapidly arise to euthermic state. Thus, during hibernation period, Tb does not remain constant at neither low nor high values but shows fluctuation between euthermia and hypothermia with an interval of several days. This multiday-scale-phenomena, known as torpor-IBA cycles, is a conserved property of hibernation across mammalian hibernators.

IBA is also called periodic arousal (Lyman et al., 1982), for patterns of Tb fluctuation in the middle of hibernation season attempted one to find certain periodicity in torpor-IBA cycles. However, length of torpor bout gradually changes in the early and late period of hibernation even under a constant condition in a laboratory, casting questions whether the torpor-IBA cycle is explicitly defined as “periodic” and what biological processes are behind it. Several hypotheses have long been proposed to explain the regulation and significance of the torpor-IBA cycles (Andrews, 2019). There are two non-mutually exclusive hypotheses currently proposed. First is that the timing of IBA is regulated by accumulation or consumption of certain proteins, protein modifications, or metabolites during torpor period. And the second one is that its timing is a reflection of certain innate endogenous rhythm(s) in the animal, such as circadian or circannual rhythms (Hampton & Andrews, 2007). In nature and human societies, some systems exhibit a gradual change in the period of oscillations (Granada et al., 2011). For instance, timing of sleep onset for some non-24 h patients with sleep-wake disorder is delayed every day and fluctuates several times a month (Uchiyama et al., 1996). Theoretically, this phenomenon can be understood as the desynchrony between circadian rhythms and 24 h environmental cycles; however, it has not been tested whether such desynchronization is responsible for producing Tb patterns during hibernation. Now that recent technology development enables to monitor Tb for more than hundred days with high precision, quantitative analysis and phenomenological modeling of Tb time-series data can be used to address the principle governing hibernation.

The frequency change in biological time series is often quantified by short-time Fourier transform (STFT) and Wavelet transform. In STFT, the time series, which is multiplied by short interval window function, is analyzed, instead of analyzing the original time series. In Wavelet transform, the basis function is localized in time and frequency, called Wavelet function, which is distinct from trigonometric function. These two methods have been widely accepted in the field of time-series analysis. However, it is often difficult for the two methods to estimate the period of the signal within short time intervals (such as analyzing the patterns of Tb fluctuation during hibernation that can be used to uncover certain periodicity in torpor-IBA cycles) because of the fundamental tradeoff between time and frequency resolutions. To estimate the period of the signal within short time intervals, Generalized Harmonic Analysis (GHA), a methodology usually applied to acoustics for characterization of irregularities in music or circadian rhythms, can be applied to Tb fluctuation during hibernation. In contrast to STFT and Wavelet transforms, GHA can be advantageous because it simply fits the data by the summation of trigonometric functions based on least square, and it is not bounded by trade-off between time and frequency resolutions.

Mammalian hibernators are roughly classified into two types: (i) obligate (or strongly seasonal) and (ii) facultative (or opportunistic) types (Geiser, 2013) (Giroud et al., 2020). Obligate hibernators, such as ground squirrels, marmots, and bears, undergo fall transition and enter hibernation spontaneously even without environmental cues (Zucker, 1985) (Körtner & Geiser, 2000) (Schwartz & Andrews, 2013). For example, 13-lined ground squirrels and chipmunks exhibit hibernation iteratively with a period of ~1 year under conditions of constant cold and continuous darkness (Kondo et al., 2006) (MacCannell & Staples, 2021). These lines of evidence suggest that endogenous circannual rhythms underlie hibernation in these species, although mechanisms for the circannual rhythms are unknown. In contrast, facultative hibernators, such as Syrian hamsters *(Mesocricetus auratus)*, enter hibernation at any time of year, at least in a laboratory condition, when they are exposed to cold and short photoperiodic conditions (Janský et al., 1984) (Chayama et al., 2016). Consistent with the successful induction of hibernation irrespective of seasons in appropriate laboratory conditions, little evidence has been found so far that circannual control of hibernation exists in this species. Nevertheless, this species awakens from hibernation without any external factors (Janský et al., 1984) (Chayama et al., 2016), implying the existence of mechanisms for estimation of the hibernation length. However, few clues about this phenomenon exist, not only at molecular level but also the underlying mathematical principle in both obligate and facultative hibernators.

In this study, we performed theoretical analysis of Tb across hibernation cycle (>50 days) in Syrian hamsters (Chayama et al., 2016). Typically, the Tb time series during hibernation is noisy and fluctuates irregularly, which complicates analysis. Therefore, to uncover temporal changes of torpor-IBA cycle, we used GHA, a methodology usually employed for acoustics for characterization of irregularity in music.

## Results

### Dominant period in body temperature fluctuation during hibernation of Syrian hamsters changes at hundred-days scale

To find rules for Tb fluctuation during hibernation, we applied GHA to analyze Tb from 25 hibernating Syrian hamsters (Chayama et al., 2016) (Figure 1A, B, Figure S1, Figure S2). GHA enables us to accurately quantify periodic components underlying several torpor-IBA cycles (Figure 1C-F) and determine the strongest periodic component (Figure 1G, H), the so-called dominant frequency (“Material and methods” (Terada et al., 1994)). Because the torpor-IBA cycles take several days (Figure 1A-C), we extracted dominant frequency of the torpor-IBA cycles ranging from 0.1 to 0.3 per day (Figure 1E, F), which corresponds to 3.3–10 days dominant period calculated as the inverse of the dominant frequency (Figure 1G, H) to reveal the temporal changes of periodic components during hibernation. The extracted dominant periods underlying Tb time series were 3.5–8.5 days during the first 16 days after the onset of first torpor bout and 4.5–8.0 days during 68–84 days. This GHA analysis quantitatively revealed that dominant period of torpor-IBA cycle gradually changed (Figure 1I, J). The dominant period increases or decreases over time during hibernation depending on the individual (Figure S3, Figure S4). To compare patterns of Tb time series, we quantified the change of torpor-IBA cycle period at the initial 0–48 days of hibernation using linear regression (“Methods”, Figure 1K, L, Figure S3, Figure. S4). For 22 of 25 individuals (except #3, 15, and 20 in Figure S3, Figure S4), the slope of regressed line was positive, and its average was 0.0068 per day, which indicates that the period of torpor-IBA cycle increased 0.68% per day on average with time during the initial 0–48 days (Figure. 1M). Particularly, for five of twenty-five individuals, who hibernated for more than 120 days (#1, 2, #8–10), the dominant period of torpor-IBA cycle initially increased and decreased later toward the end of hibernation (Figure 1G-L, Figure S3I, L-N, P, Q). Although three of twenty-five individuals (#3, 15, and 20) exhibited strongly fluctuated Tb pattern and its slope of the linear regression was negative, it was due to the occurrence of shallow/daily torpor and long IBA during first 20 days of hibernation because the dominant period of torpor-IBA cycle exhibited an increasing tendency thereafter (see Figure S1A, Figure S2B, M, Figure S3A, D, Figure S4B, E, M, P for details). Taken together, this analysis demonstrated that the period of torpor-IBA cycle changes at hundreds-days scale, suggesting that it is possibly governed by as-yet-unknown physiological temporal process.

**Figure 1.**
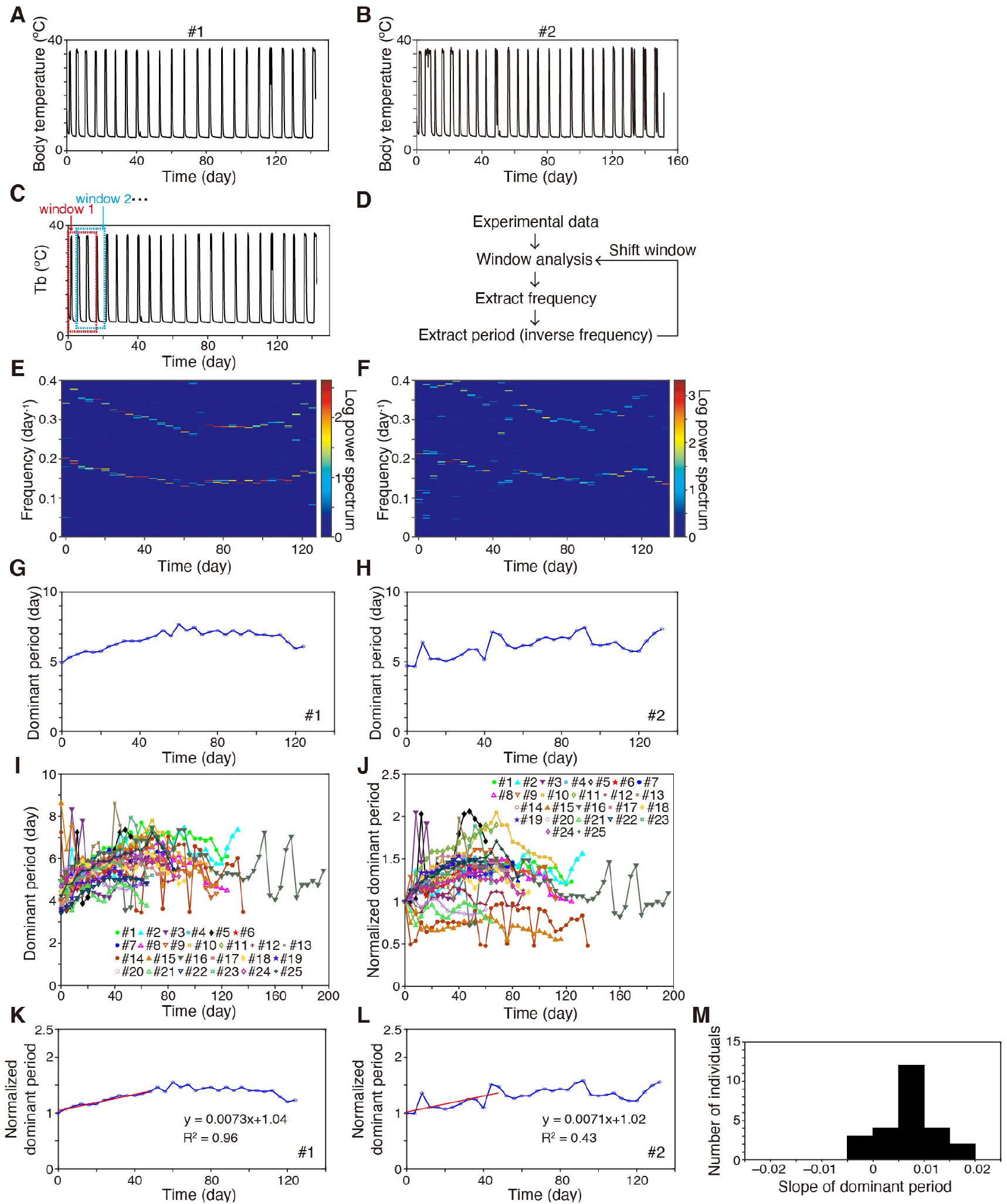
Dominant period in body temperature fluctuation during hibernation of Syrian hamsters changes at hundred-days scale. (**A, B**) Time series of Tb fluctuation during hibernation under a short day and cold temperature (8L:16D cycle, ambient temperature = 4 °C). Two representative hibernating Syrian hamsters (from n = 25) are shown in Fig. 1A, B (see also Fig. S1, Fig. S2 for data of the others). (**C, D**) Scheme of GHA analysis for Tb data (see text). (**e, f**) Sequence of estimated frequency by analysis of data from two representative individuals during hibernation (#1, 2). The heatmaps show the magnitude of spectrogram as logarithmic compression of power defined by log (1 + |amplitude|^2^). (**G-I**) Estimated dominant period (i.e., day/frequency) for #1 (**G**), #2 (**H**), and all 25 individual hamsters (**I**) changed over time. (**J-L**) Dominant periods normalized by the initial dominant period. (**K, L**) The change of dominant period for individuals #1 and #2 at 0–48 days was quantified using liner regression. (**M**) Distribution of quantified slopes of regression line per day for the change in normalized dominant period.

### Determination of a model reproducing the pattern of torpor-arousal cycles

To understand biological processes behind gradual change of the torpor-IBA cycle, we tested two theoretical models that possibly reproduce the experimental data, (a) frequency modulation model (FM) and (b) desynchrony model (Figure 2A, B). FM assumes that the shorter frequency (period) may be modulated by another slower frequency (longer period). Its typical example is FM radio exhibiting gradual change in period over time^17^, although few biological processes other than auditory and vocalization systems were proposed to exhibit such features. In contrast, the desynchrony model assumes that one period, torpor-IBA cycle period in this case, changes over time due to the failure of synchronization with another period. A typical example of this is desynchronization of internal circadian rhythm and external photoperiod (Figure 2A, B, “Methods”).

**Figure 2.**
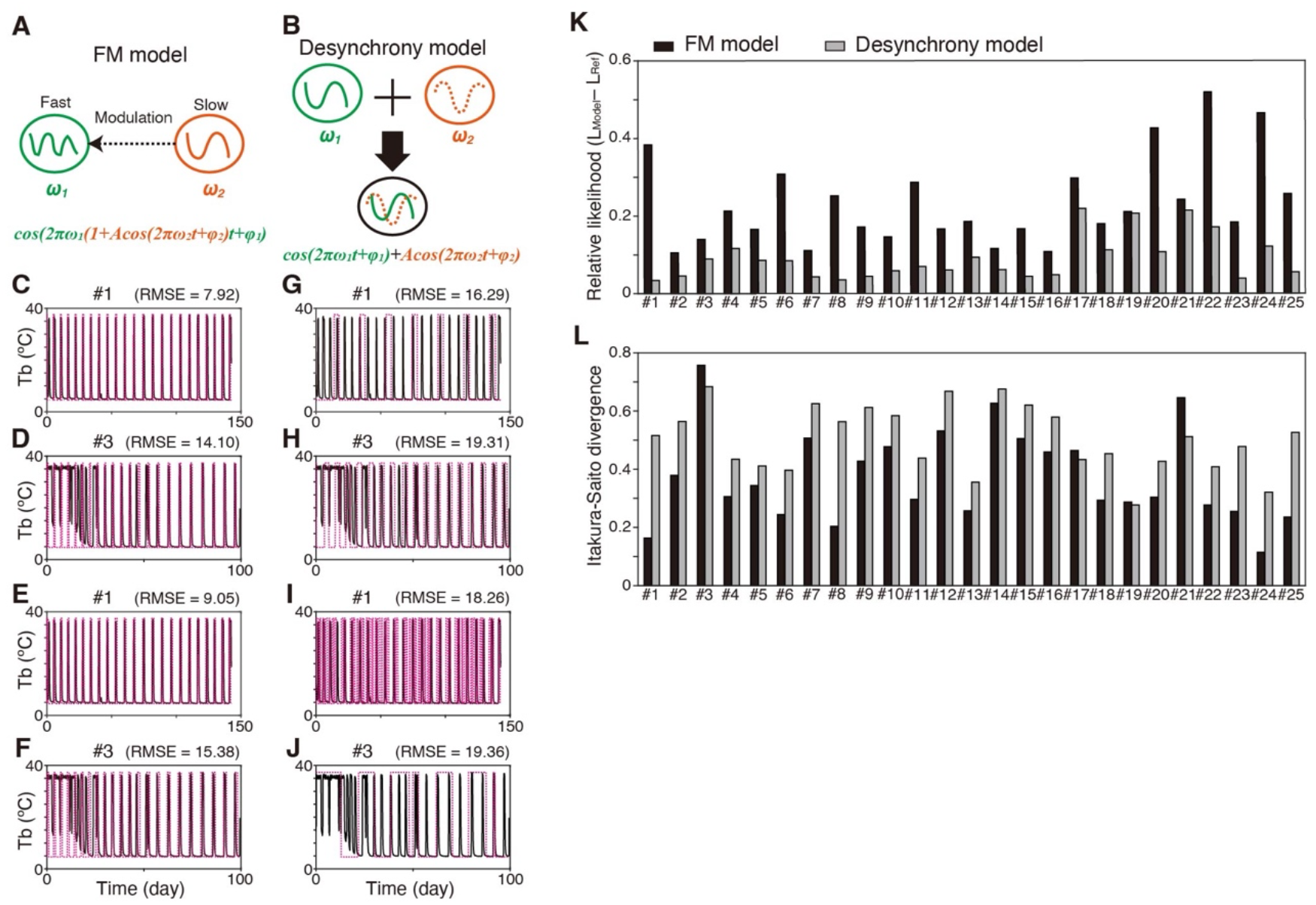
Determination of a model reproducing the pattern of torpor-arousal cycles. (**A, B**) Schematic representation of the proposed FM model (**A**) and desynchrony model (**B**) with multiple frequencies of *ω*_1_ and *ω*_2_. (**C-J**) Comparison of simulation done by the two models. (**C-F**) FM model simulation (red) with the best parameter set for Tb time series recorded in two representative animals (black, #1,3). The best parameter was chosen by maximum likelihood estimation (**C, D**) and IS divergence (**E, F**). RMSE is the value of root mean squared error. (**G-J**) Desynchrony model simulation (red) with the best parameter set for the Tb data from the same animals used for FM simulation (#1,3). The best parameter was chosen by maximum likelihood estimation (**G, H**) or IS divergence (**I, J**). (**K, L**) Maximum likelihood estimation comparison of the two models to realize each individual time series (#1– 25). Likelihood of FM (black) and desynchrony model (gray) was compared using maximum likelihood estimation (**K**) and IS divergence (**L**). Time series of fixed minimum of Tb was set as the reference model. Note that as the model becomes closer to the experimental data, maximum likelihood estimation becomes larger and IS divergence becomes smaller.

By varying the parameters in both models, we first investigated which model would better reproduce the observed Tb fluctuation using maximum likelihood estimation. The FM model with the best parameter set, yielding maximum likelihood, reproduced most of the timing for the transitions between deep torpor and IBA for all individual data (Figure 2C, D). In contrast, the desynchrony model did not reproduce most transitions between deep torpor and IBA for some individuals (Figure 2G, H). Indeed, likelihood values of the FM model with the best parameter set were always larger for each individual’s data than those of the desynchrony model, suggesting that the FM model is the more proper model for reproducing Tb fluctuation of Syrian hamsters during hibernation (Figure 2K, Figure S5-8 in detail).

We next evaluated the agreement between theoretical models and Tb experimental data by other statistical method, the so-called “Itakura–Saito (IS) divergence.” IS divergence may be a better statistical method to identify the model and its parameter set that reproduces the timing of IBA than widely-used maximum likelihood estimation (Itakura & Saito, 1968) from the following points; IS divergence, originally a speech recognition method for mixed sound, has axial asymmetry while the Euclidean distance has axial symmetry. Thus, IS divergence imposes more penalties if the value of the model is smaller than that of experimental data (Hashimoto et al., 2014). Indeed, FM model with the best parameter set, yielding minimum IS divergence, reproduced most of the timing of IBAs (Figure 2E, F, Figure S9, 10). Quantification of the maximum likelihood estimation using IS divergence demonstrated that the FM model is much better than the desynchrony model in reproducing Tb fluctuation (Figure 2L, for examples compare #5, #8, and #12 in Figure S9). For three individuals wherein the desynchrony model showed a better score than the FM model (#3, 17, 21 in Figure 2L), it was evident that FM model fits the Tb data much better than desynchrony model for the most part in the time series (Figure 2F, J, Figure S10G, H, O, P), suggesting that the judgment with IS divergence is inappropriate for the individual. Taken together, both maximum likelihood estimation and IS divergence statistically justified that the FM model is a proper model for reproducing periodic change of torpor-IBA cycles.

### Two endogenous periods, several days and hundreds of days, underlie hibernation

The above result indicates that the main property of Tb fluctuation during hibernation can be described by the two key parameters in the FM model, shorter (day/*ω*_1_ and longer (day/*ω*_2_) period. In this model, shorter period is modulated by longer period (>*ω*_2_). By determining the two periods with the theoretical model, we can now quantify and explain individual variation in complex patterns of Tb fluctuation. Because the dominant period estimated by GHA was 3.5–9 days and the longer period at the timescale of approximately 100–500 days, we first rigorously varied the shorter period at the timescale of approximately 1–10 days to precisely determine the two periods (see “Methods”). To this end, we compared estimations by maximum likelihood and IS divergence as follows.

Based on the best maximum likelihood estimation values for the FM model that reproduced experimental data (Figure 3A-C, Figure S11, Figure S12), we determined the faster and slower period (*ω*_1_, and *ω*_2_); the shorter period of the FM model, yielding the maximum likelihood estimation, was between 3.5 and 9.2 days, which covers the dominant period extracted by GHA (Figure 3D). Meanwhile, the estimated longer period was between 119 and 430 days (Figure 3D). Conversely, when the parameter set of the FM model was chosen by IS divergence, the distribution of the shorter period of the FM model (day/*ω*_j_ was narrower such that it was between 3.5 and 6.5 days (Figure 3E). The longer period by IS divergence (day/*ω*_2_) was also between 115 and 430 days (Figure 3E). Therefore, there was a discrepancy between the maximum likelihood and IS divergence estimation. It was probably because the FM model yielding the maximum likelihood estimation did not reproduce shallow/daily torpor of some individuals, whereas the FM model yielding minimum IS divergence did reproduce them (see #5 and #7 in Figure S5, Figure S9). Taken together, analysis with the FM model enabled us to quantify a period of several days and another longer period of 115–430 days that modulates the former period during hibernation.

**Figure 3.**
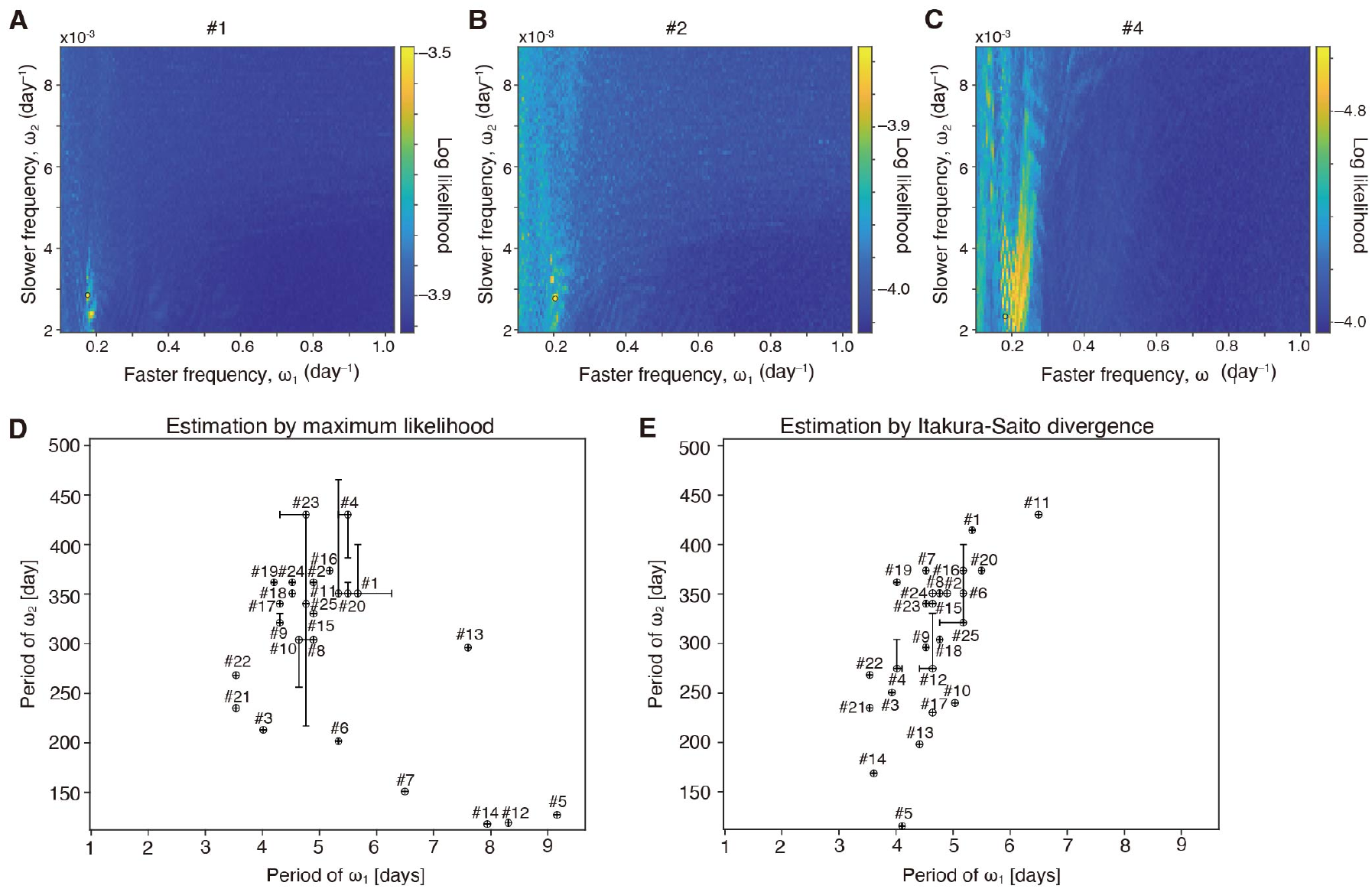
Two endogenous periods, several days and hundreds of days, underlie hibernation. (**A-C**) Distribution of likelihood values for the set of faster and slower period (*ω*_1_ and *ω*_2_ in FM model) underlying Tb fluctuation was estimated by maximum likelihood. Individual variation in the set of faster and slower period estimated using maximum likelihood (**D**) and IS divergence (**E**).

### Forecasting body temperature fluctuation during hibernation

The validity of the FM model for Tb fluctuation can be assessed by its predictability. To test this, we estimated the two parameters (day/and *ω*_2_) from three-quarters (Figure. 4A, B) of the whole time series of Tb data for each individual’s data by the FM model using maximum likelihood (Figure. S13, Figure. S14) or IS divergence (Figure. 4, Figure. S15, Figure. S16). Simulation with the parameters predicted the last quarter of the time series because the estimated shorter and longer period (day/and day/*ω*_2_) from the three-quarters of the Tb time series were close to those estimated from the whole time series (Figure. 4A, B). When we simulated the FM model with the parameters estimated from the first half of the whole time series of the Tb data, it also predicted the first few cycles of the second half of the original whole Tb time series but gradually deviated from the original (Figure. 4C, D). Indeed, the longer periods (day/*ω*_2_) predicted from the first half were distant from those estimated from the original whole Tb time series (Figure. 4E, G), implying that the estimation of the parameter needs sufficiently long experimental Tb time series (Figure. 4E, G). In contrast, when the shorter frequency (ωj was predicted from the first half of the whole time-series data, it ranged from 3.4 to 5.9 days and was close to that estimated from the whole data (Figure. 4E, F). These results suggest that estimation of longer frequency (*ω*_2_) is affected by and that of shorter frequency (*ω*_1_ and is robust to the length of Tb time series used for statistical analysis.

**Figure 4.**
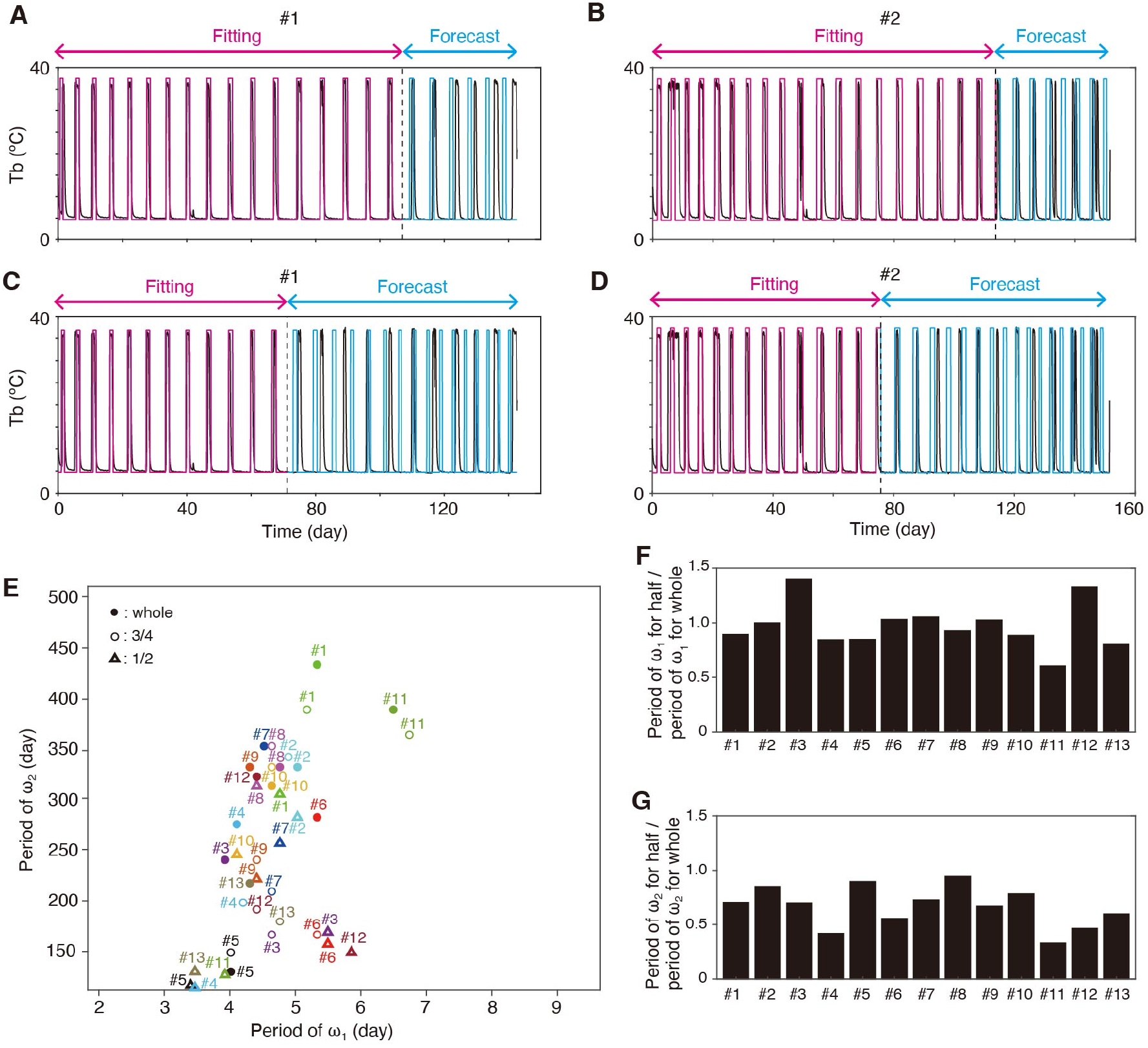
Forecasting body temperature fluctuation during hibernation. Tb time series was reconstructed (magenta) and predicted (cyan) using the three-quarters (**A, B**) and the first half (**C, D**) of the whole time series of original Tb data in animals (black). (**E**) Predicted faster and slower period (*ω*_1_ and *ω*_2_ in FM model) using partial time series based on IS divergence estimation. (**F**) The ratio of *ω*_1_ period for half time series to that for whole time series. (**G**) The ratio of *ω*_2_ period for half time series to that for whole time series.

## Discussion

In this study, we applied GHA to quantify the time series of Tb fluctuation during hibernation and defined theoretical models that reproduce it. This study demonstrates the effectiveness of GHA to quantify Tb fluctuation, the patterns of which are complex and rich in variation among animals, and is an improvement over using STFT and Wavelet transform to explore and model Tb oscillations.

Through statistical analysis of the FM model and desynchrony model, we conclude that the FM model is better than the desynchrony model to realize the pattern of Tb fluctuation during hibernation of Syrian hamsters. To the best of our knowledge, only a few studies have applied FM to explain biological processes, such as circadian desynchronization in rats and changes in oscillatory frequency of brain waves in visual perception (Granada et al., 2011) (Wutz et al., 2018). The FM model hypothesizes that the frequency of oscillation (*ω*_1_) gradually changes over time with another longer frequency (*ω*_2_). Our statistical analysis derives realistic values for *ω*_1_ and *ω*_2_. The derived ranges of both values in the case of analysis with IS divergence were 3.9–6.5 day period for *ω*_1_ and 115–430 day period for *ω*_2_, well within the 2–3 year lifespan of Syrian hamsters. This result raises the question of what biological processes are reflected in these values.

The faster frequency *ω*_1_, a period with a few days, could be responsible for determining the period of torpor-IBA cycles. In many small hibernators, length of each torpor bout gradually increases from the beginning, reaches its peak in the middle, and then decreases at the end of the hibernation season, even in constant ambient temperature conditions (MacCannell & Staples, 2021) (Ortmann & Heldmaier, 2000) (Arnold et al., 2011) (Sheriff et al., 2013) (Siutz et al., 2018). Ambient and core body temperatures also affect duration of each deep torpor bout, suggesting involvement of a temperature-sensitive process in the regulation of torpor-IBA cycles (MacCannell & Staples, 2021) (Twente & Twente, 1965) (Geiser & Kenagy, 1988) (Malan et al., 2018). Metabolic rate measured by oxygen consumption contributes to the determination of torpor-IBA timings in golden mantled ground squirrels and garden dormice (Geiser & Kenagy, 1988) (Ruf et al., 2021), although the exact metabolic process responsible for determination of torpor-IBA timing is not yet clear. Most chemical reactions are temperature-sensitive. It should be noted that the temperature-sensitive nature of torpor-IBA cycle is apparently different from temperature-compensation of circadian rhythm (Pittendrigh, 1954) (Kurosawa et al., 2017). In fact, several studies suggested that clock genes involved in circadian rhythm have little or small contribution to determining timing of deep torpor-IBA cycles during hibernation (Oklejewicz et al., 2001) (van der Vinne et al., 2018). These lines of evidence suggest that mechanisms other than circadian rhythm governed by transcription-translation feedback loop underlying the timing of torpor onset and arousal during hibernation. *ω*_1_ may correspond to the period generated by such mechanisms. In contrast, timing of shallow/daily torpor, an adaptive response to winter condition or food shortage, during which basal metabolisms and Tb drops for a shorter period, typically several hours and within a day, is related to circadian rhythm in some species, including Djungarian hamster (*Phodopus sungorus*), which exhibits seasonal shallow/daily torpor, and mice entering fasting-induced torpor (Körtner & Geiser, 2000) (Malan et al., 2018) (Oklejewicz et al., 2001) (van der Vinne et al., 2018) (Kirsch et al., 1991) (Ruf & Geiser, 2015). Thus, the situation is still complex and further research is needed to determine whether is sensitive to ambient temperature during hibernation.

Another frequency *ω*_2_, a period of a few hundred days, implies endogenous circannual rhythm. Although we recognize the limitation of our study that our model does not integrate the timing of the start and end of the hibernation period, it is surprising that only Tb time-series data during hibernation period could derive such a long period. In some species including mammalian hibernators and migratory birds, the circannual rhythm is regulated by unknown endogenous mechanisms independent of environmental triggers, such as photoperiod and ambient temperature (MacCannell & Staples, 2021) (Gwinner, 2003). Ground squirrels kept in a constant condition change their body weight and exhibit torpor phenotypes in a circannual cycle (MacCannell & Staples, 2021) (Mrosovsky, 1971) (Pengelley et al., 1978). Additionally, recordings over several years in ground squirrels or eastern chipmunks kept under constant photoperiods and cold temperatures demonstrated that the periods from the onset of hibernation to that in next year range from 5 months to 16 months or 5 months to 13 months in those animals, respectively (Kondo et al., 2006) (MacCannell & Staples, 2021) (Mrosovsky, 1971) (Pengelley et al., 1978). Thus, the endogenous period underlying circannual phenomena could vary among individuals, implying that coordination of the endogenous circannual period with phenological changes would be necessary for proper adaptation to animal habitat. Although Syrian hamsters are facultative hibernators that do not depend on or may not have an endogenous circannual rhythm, they spontaneously quit hibernation after several months of hibernation period under a constant cold and photoperiodic condition. This strongly suggests that Syrian hamsters have an unknown endogenous timer for measuring the length of hibernation. In fact, a phenomenon called refractoriness has long been known in Siberian and Syrian hamsters (Johnston et al., 2003) (Herwig et al., 2013) (Carr et al., 2003). The hamsters regress or regrow their gonads in response to chronic short or long photoperiod, respectively. But if the animals were kept further in short or long photoperiod conditions for several months after the gonadal regression, the gonads then start to regrow or regress without any environmental changes, suggesting the existence of yet-unidentified endogenous timer measuring seasonal length in hamsters. Thus, *ω*_2_ may correspond to possible endogenous period governed by such an endogenous timer. Interestingly, we found no significant correlation between *ω*_2_ and the hibernation duration (Figure. S17), which remind us the result that there is no correlation between the circannual period and the hibernation duration in chipmunk, an obligate hibernator (Kondo et al., 2006) (Kondo, 2007).

We anticipate that our theoretical analysis of Tb fluctuation can be a starting point for quantitative comparison of hibernation patterns across close and distantly related hibernating species exhibiting various Tb patterns. Furthermore, quantification across two or more consecutive hibernation seasons may allow prediction of the timing of torpor and IBA and will also foster the understanding of molecular mechanism of hibernation by searching for biological processes that operate within those periods.

## Material and methods

### Animal housing and Tb measurement

Animal housing and Tb measurement were done as described previously (Chayama et al., 2016). Briefly, female Syrian hamsters (*Mesocricetus auratus*) were purchased from SLC, Inc., Japan and reared under LD-Warm conditions (light condition = 16L:8D cycle, lights on 05:00–21:00, ambient temperature = 24–25°C) until most animals weighed over 100–120 g. Then the animals were subjected to surgical operation under inhalation anesthesia with 4% isoflurane (DS Pharma Animal Health, Japan) and intraperitoneal injection of pentobarbital sodium (65 mg/kg, diluted with phosphate-buffered saline; Kyoritu Seiyaku, Japan) for intraperitoneally implanting core body temperature (Tb) loggers (iButton^®^, Maxim Integrated, USA, #DS1992 L-F5 model) coated with rubber (Plasti Dip, Performix^®^). After one to two weeks of recovery, animals were transferred to SD-Cold conditions (8L:16D cycle, lights on 10:00–18:00, ambient temperature = 5°C) for hibernation induction. Animals were individually housed in polypropylene cages, and the Tb of animals were measured every 90 min with an accuracy of 0.5°C. The cage replacement was done every two weeks and skipped when animals were hibernating at deep torpor to avoid disturbing it. The Tb loggers were recovered from animals sacrificed by decapitation after they were subjected to 10–15 min anesthesia with intraperitoneal injection of pentobarbital sodium (65 mg/kg) and inhalation of 4% isoflurane.

### Proposed theoretical models for Tb fluctuation

To simulate Tb fluctuation, we tested two models, frequency modulation and desynchrony models. In the FM model, Tb during hibernation is expressed by the following equation:

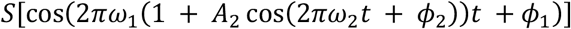

wherein oscillation with frequency of oscillation (*ω*_1_) is assumed to change over time with frequency (*ω*_2_). (b) In the desynchrony model, Tb is expressed as

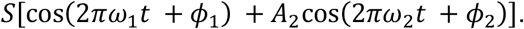

Although the desynchrony model can yield synchronous limit cycle oscillation when *ω*_1_ = *ω*_2_, it also yields quasiperiodic oscillations for certain parameter choices with *ω*_1_ ≉ *ω*_2_. Step function *S* was used in both models, which realize sharp change in Tb time series. Function *S*[*x*] is defined as *S*[*x*] = maximum value of Tb if *x* > *θ* and *S*[*x*] = minimum value of Tb otherwise. In search of the models and parameters that reproduce each data, all the parameters were rigorously varied (variation is detailed in Supplementary Tables 1, 2).

### Quantification of body temperature fluctuation

GHA was conducted as previously described for the studies of music and circadian rhythms (Terada et al., 1994) (Gibo & Kurosawa, 2020). GHA is a method in signal processing to quantify periodic components within a certain frequency range, so-called “dominant period” by estimating the most fitted summation of trigonometric functions for given time series (Terada et al., 1994). The changes of torpor-IBA period over time were quantified by [1] setting a time window; [2] estimating dominant torpor frequency of each time window by using GHA; [3] shifting time window by 4 days; and [4] repeating [2] and [3] until the end of time series. Here, the time-window length was set to be 16 days so that it is sufficient for covering 2–3 cycles of DT-PA cycles. In the experiment, 25 individual Syrian golden hamsters continued to hibernate under laboratory conditions for more than 50 days. In this analysis, the onset of hibernation is defined to be the point such that Tb is lower than 15 °C. Time series of body temperature within the time window of 16 days for the 25 hamsters is modeled as

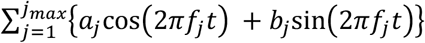

where *f_j_* is the frequency and *a_j_* and *b_j_* are the amplitudes. The resolution of frequency was 1/500 per day, and *j*_max_ is the maximum number of frequency component, which was set to be 4000 (i.e., *f_jmax_* = 8 /day). The frequency, called dominant frequency (text) and the amplitude within each time window were quantified by repeatedly minimizing the square residuals:

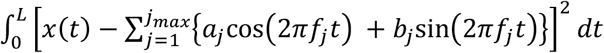

where *L* is time-window length. Then, the estimated dominant period during hibernation, which is the inverse of the dominant frequency was plotted over time (Figure 1).

After quantifying changes in the dominant period of torpor-IBA cycle during hibernation, the change in the dominant period during the initial 0–48 days of hibernation was specifically measured by linear regression. The positive (negative) slope of the regression line indicates that the period of torpor-IBA cycle increases (decreases) with time.

### Statistical analysis for model and parameter selection

In search of theoretical models and their parameters reproducing the experimental Tb data during hibernation, we used the following two statistical quantities: maximum likelihood and Itakura–Saito divergence (IS), defined as follows, respectively:

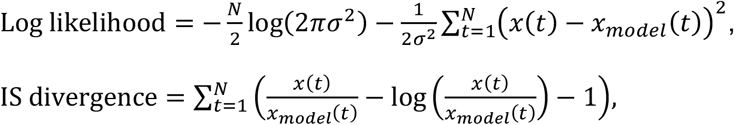

where *x*(*t*) is original time series, *X*_model_(*t*) is time series of model, *N* is length of time series, and *σ*^2^ is variance of *x*(*t*)-*x*_model_(*t*). Analysis using IS divergence was conducted as previously described in the field of speech recognition (Itakura & Saito, 1968). In fact, IS divergence is small when the model reproduces the timing of periodic arousal of experimental data.

To find the best model and the best parameter for each model to reproduce the experimental data, we varied parameters as shown in Supplementary Tables 1 and 2. When maximum likelihood becomes larger, IS divergence becomes smaller; the model becomes closer to the experimental data. The best parameter for each model to reproduce the experimental data was computationally identified. Maximum likelihood and IS divergence of each model using the best parameter is shown for each individual’s data in Figure. 2K, L. There was a discrepancy between maximum likelihood and IS divergence estimation. This is because maximum likelihood often imposes more penalties than IS divergence does if the difference of *x*(*t*) and *X*_model_(*t*) is large; maximum likelihood assumes that the difference of *x*(*t*) and *X*_model_(*t*) follows normal distribution while IS divergence assumes that it follows asymmetrical distribution.

To obtain the precise values of and *ω*_2_ in FM model, the best parameter was searched within a narrow range of and *ω*_2_, after it was searched within various parameter values (Table 1). Also, in the analysis of prediction of Tb fluctuation (Figure 4), the lower limit of was set to be 0.5358 /days (1.83 days) for computation cost because the likelihood of the model with the period around 24h was always small.

## Acknowledgements

We thank Y. Chayama and H. Taii for their help in collecting Tb data; R. Enoki, E. Gracheva, and Y. Kawahara for comments on the manuscript; and A. Mochizuki for encouragement. We would like to thank Enago for the English language review. This work was supported by grants from Japan Science and Technology Agency (JPMJCR1913 to G.K.), and from the Japanese Society for the Promotion of Science, and the Ministry of Education, Culture, Sports, Science, and Technology in Japan (JP20H05766 to Y.Y. and JP21K06105 to G.K.), Toray Science Foundation (to Y.Y.), the Takeda Science Foundation (to Y.Y.), and Inamori Foundation (to Y.Y.).

## Competing interests

No competing interests declared.

## Data availability

All original data are available upon reasonable request to the corresponding authors Y.Y and G.K.

## Notes

### Competing Interest Statement

The authors have declared no competing interest.

### Summary of Updates

Supplemental file updated.

