## Supplementary file for "Frequency modulated timer regulates mammalian hibernation"

Shingo Gibo[1], Yoshifumi Yamaguchi[2,3,4,*], and Gen Kurosawa[1 *]
[1]RIKEN Interdisciplinary Theoretical and Mathematical Sciences Program (iTHEMS); Wako, 351-0198, Japan
[2]Institute of Low Temperature Science, Hokkaido University; Kita-19, Nishi-8, Kita-ku, Sapporo, 060-0819, Japan
[3]Global Station for Biosurfaces and Drug Discovery, Global Institution for Collaborative Research and Education (GI-CoRE), Hokkaido University; Kita-12, Nishi-6, Kita-ku, Sapporo 060-0812, Japan.
[4]Inamori Research Institute for Science Fellowship (InaRIS); 620 Suiginya-cho, Shimogyo-ku, Kyoto 600-8411, Japan.

**This PDF file includes:**

Figures S1 to S17
Tables S1 to S2

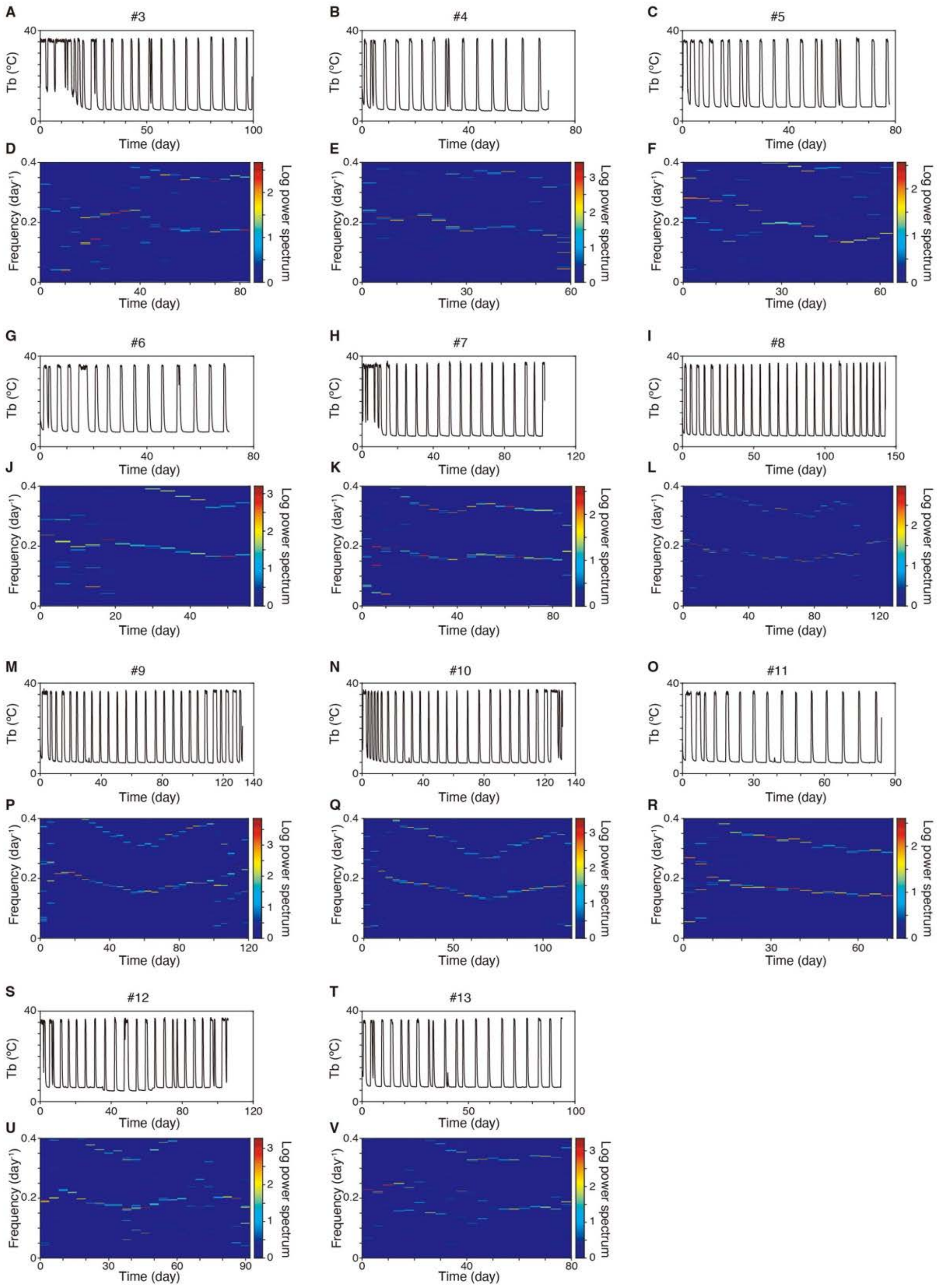


**Fig. S1. Magnitude of spectrogram by the GHA analysis of Tb data during hibernation (A-V).** Spectrogram as logarithmic compression of power (log(1+|amplitude|$^2$)) is plotted. In our analysis, the onset of hibernation is defined to be the point such that Tb is lower than 15 degree. Animal ID (#3-13) are shown on the top of graphs. Total 25 individual data including 2 in Fig. 1A, B, E, F and 12 in Fig. S2 were analyzed.

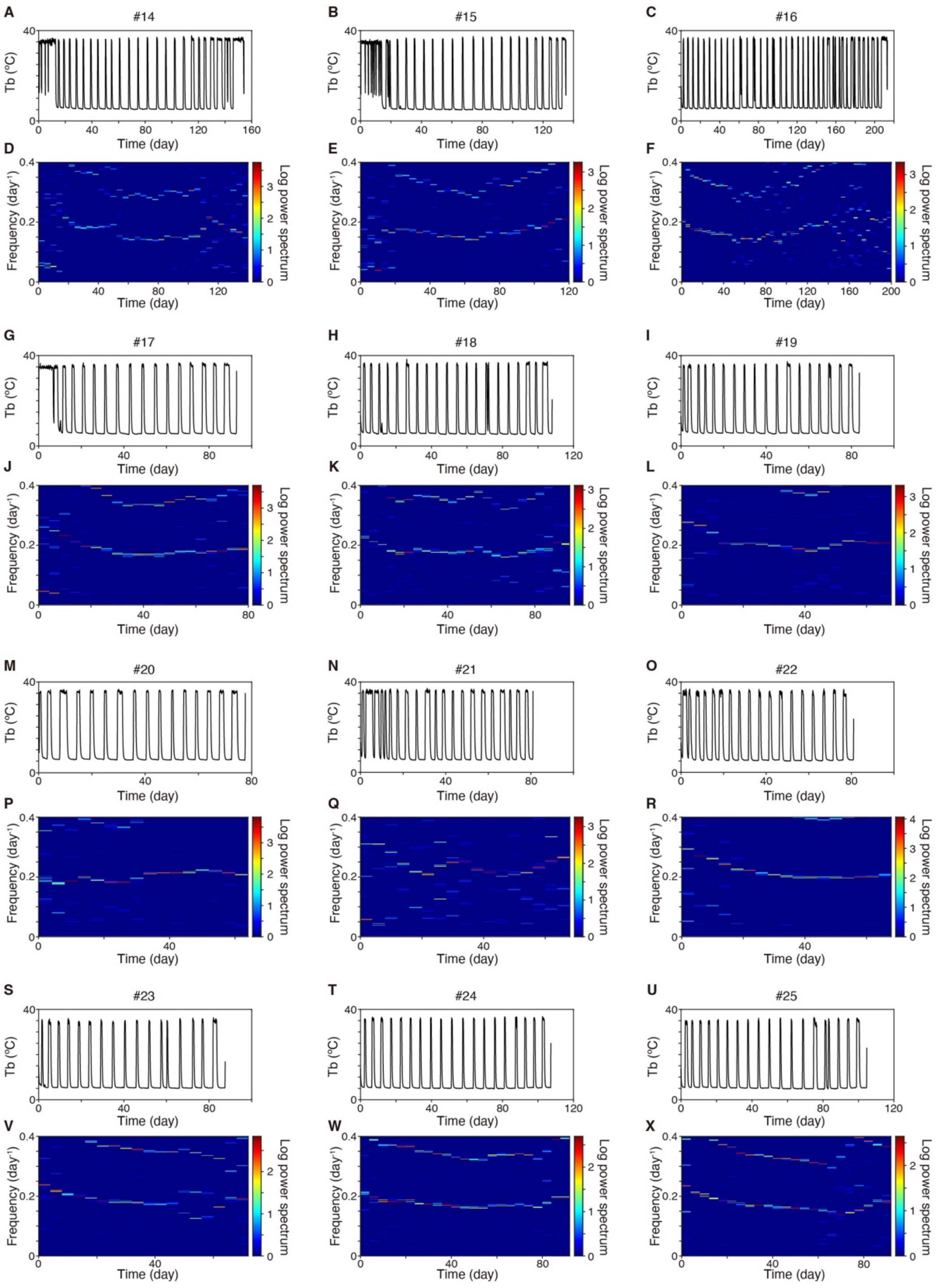


**Fig. S2. Magnitude of spectrogram by the GHA analysis of Tb data during hibernation (A-X).** Spectrogram as logarithmic compression of power (log(1+|amplitude|$^2$)) is plotted. In our analysis, the onset of hibernation is defined to be the point such that Tb is lower than 15 degree. Animal ID (#14-25) are shown on the top of graphs. Total 25 individual data including 2 in Fig. 1A, B, E, F and 11 in Fig. S1 were analyzed.

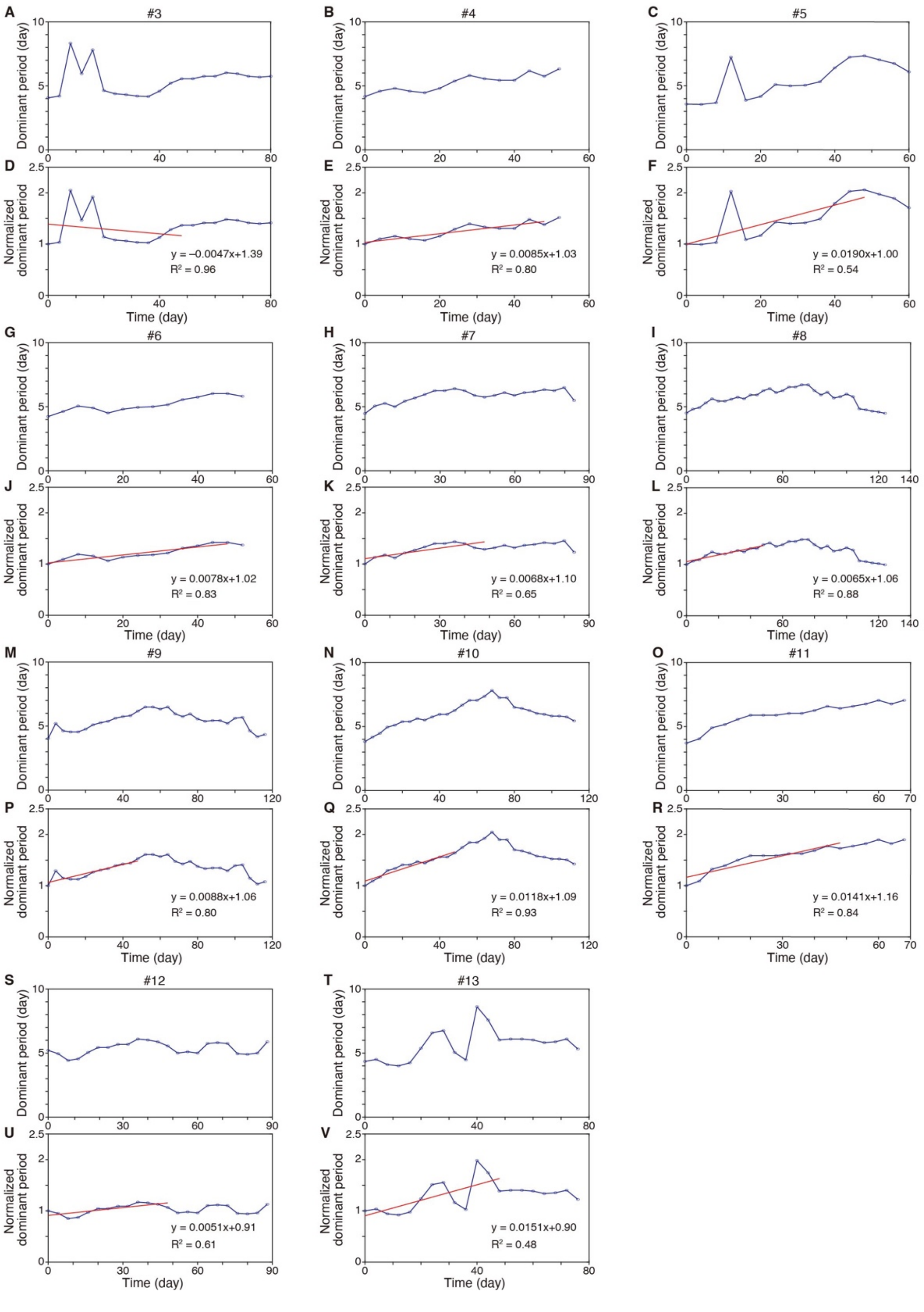


**Fig. S3. Changes in dominant period (i.e. 1/frequency) over time, estimated by the GHA analysis (A-V).** Dominant period was normalized by using the initial dominant period. Red line is regression line for the normalized dominant period at the 0-48 days. Animal ID (#3-13) is shown on the top of each graph. Total 25 individual data including 2 in Fig. 1G, H, K, L and 12 in Fig. S4 were analyzed.

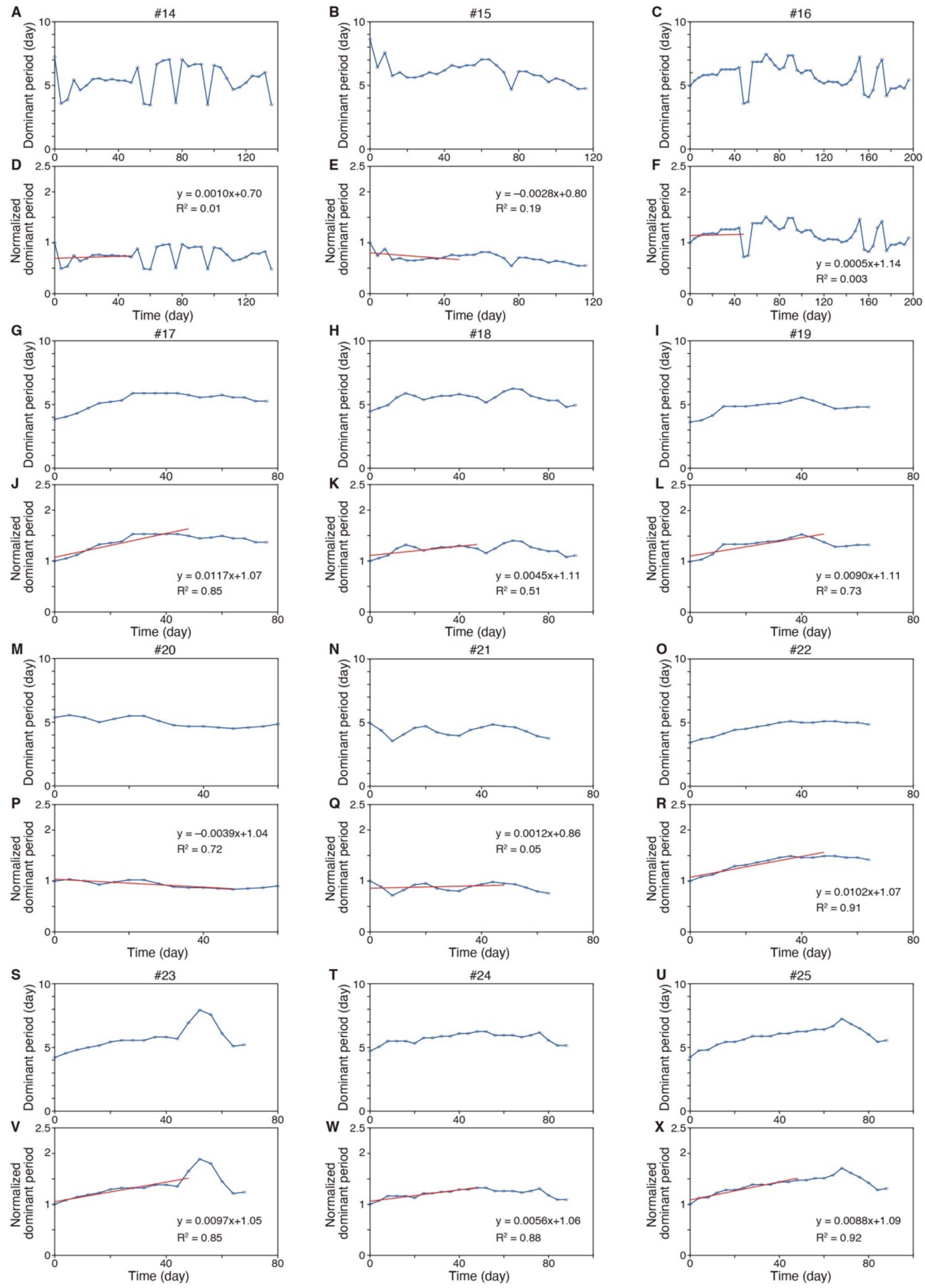


**Fig. S4. Changes in dominant period (i.e. 1/frequency) over time, estimated by the GHA analysis (A-V).** Dominant period was normalized by using the initial dominant period. Red line is regression line for the normalized dominant period at the 0-48 days. Animal ID (#14-25) is shown on the top of each graph. Total 25 individual data including 2 in Fig. 1G, H, K, L and 11 in Fig. S3 were analyzed.

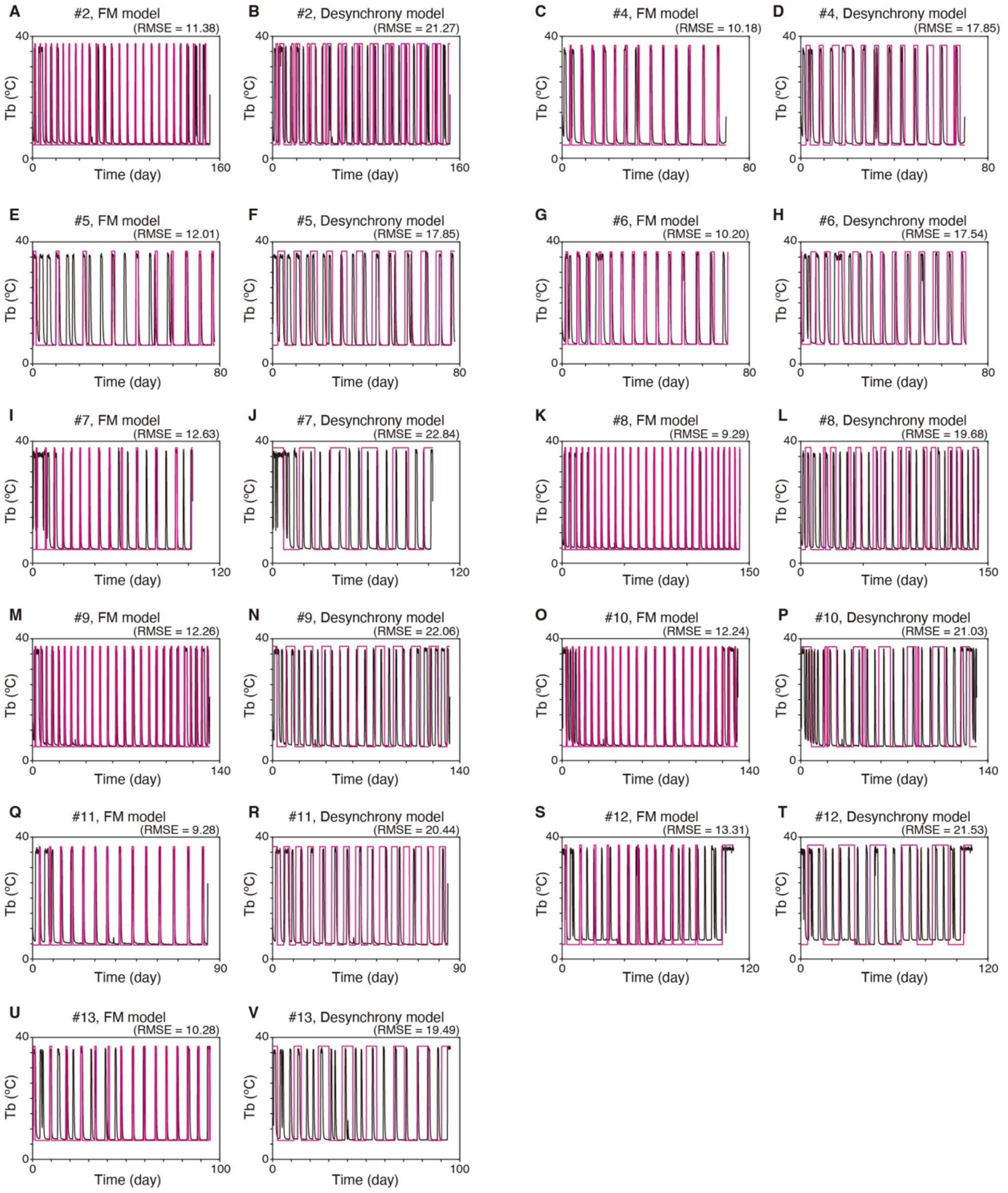


**Fig. S5. Frequency modulation and desynchrony model based time-evolution (red) best-fitted for recorded Tb data (black) of 11 animals (A-V).** The best-fitted parameter was chosen using maximum likelihood. Animal ID (#2, 4-13) is shown on the top of each graph. RMSE is the value of root mean squared error. Total 25 individual data including 2 in Fig. 2C, D, G, H and 12 in Fig. S6 were analyzed.

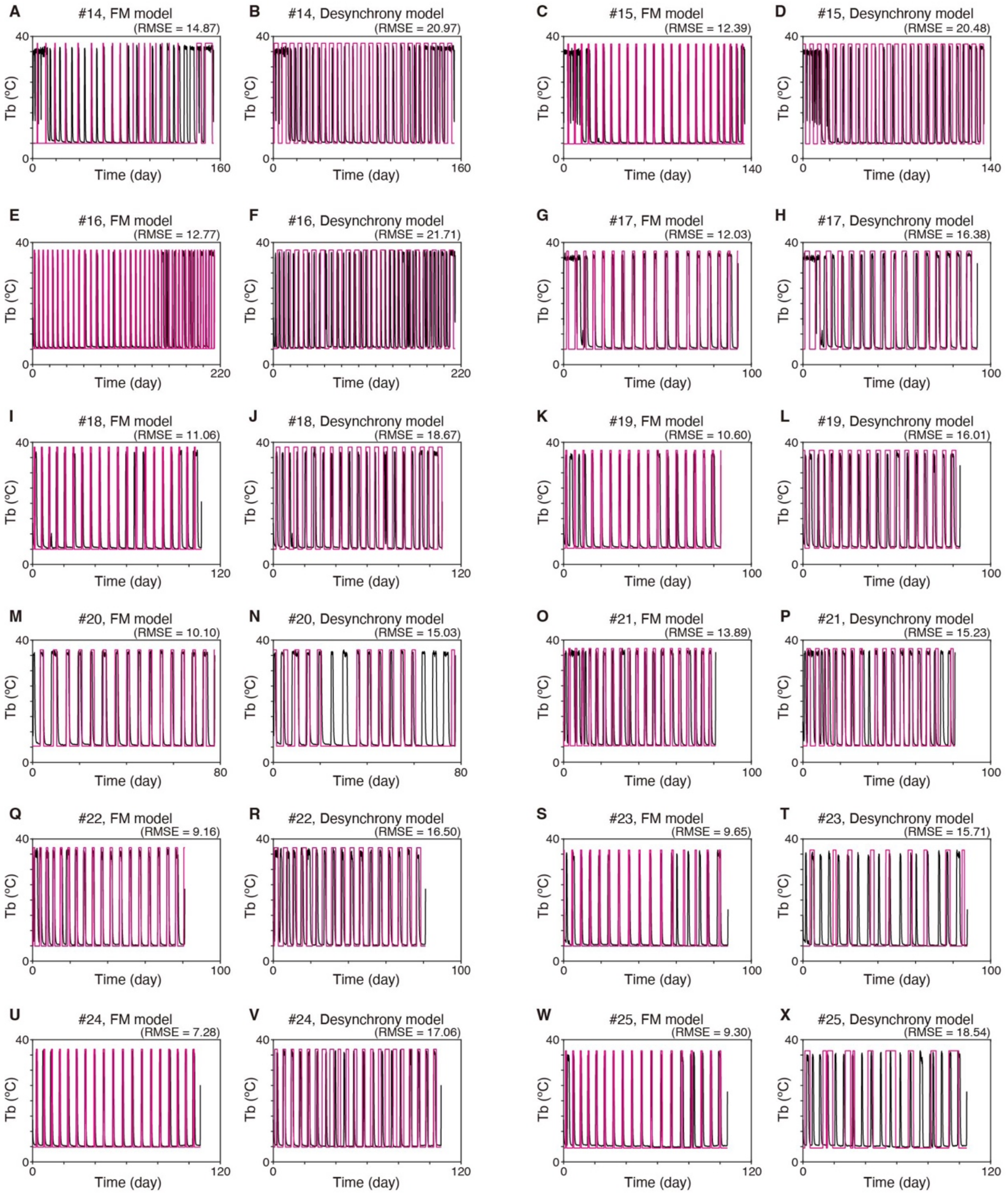


**Fig. S6. Frequency modulation and desynchrony model based time-evolution (red) best-fitted for recorded Tb data (black) of 12 animals (A-V).** The best-fitted parameter was chosen using maximum likelihood. Animal ID (#14-25) is shown on the top of each graph. RMSE is the value of root mean squared error. Total 25 individual data including 2 in Fig. 2C, D, G, H and 11 in Fig. S5 were analyzed.

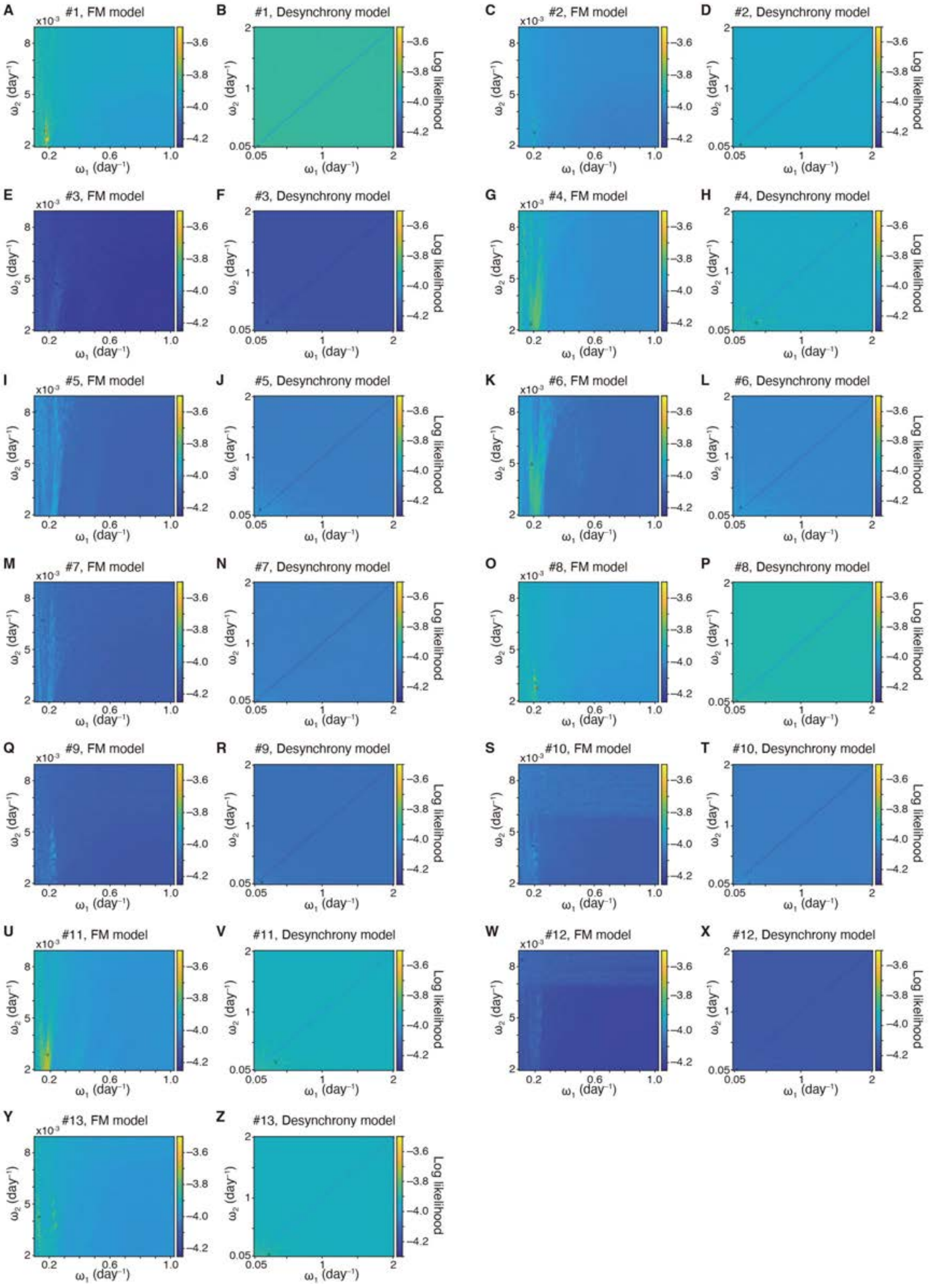


**Fig. S7. Distribution of log likelihood as a function of two frequencies ($\omega_1$ and $\omega_2$) of FM and desynchrony model for 13 individual experimental data (A-Z).** The best-fitted parameter, yielding the maximum likelihood (circle) was used in Fig. 2C, D, G, H, Fig. S5.

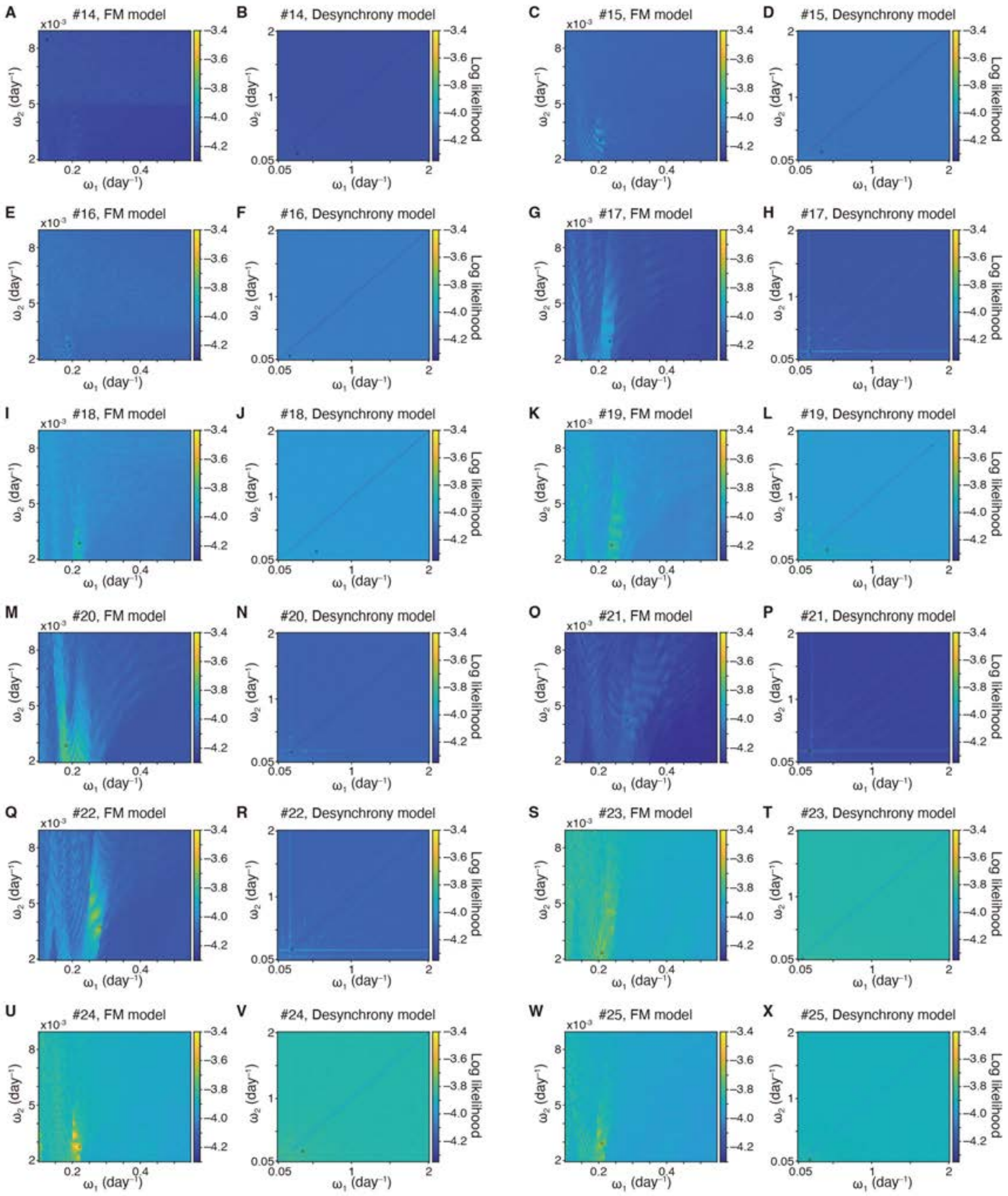


**Fig. S8. Distribution of log likelihood as a function of two frequencies ($\omega_1$ and $\omega_2$) of FM and desynchrony model for 12 individual experimental data (A-Z).** The best-fitted parameter, yielding the maximum likelihood (circle) was used in Fig. S6.

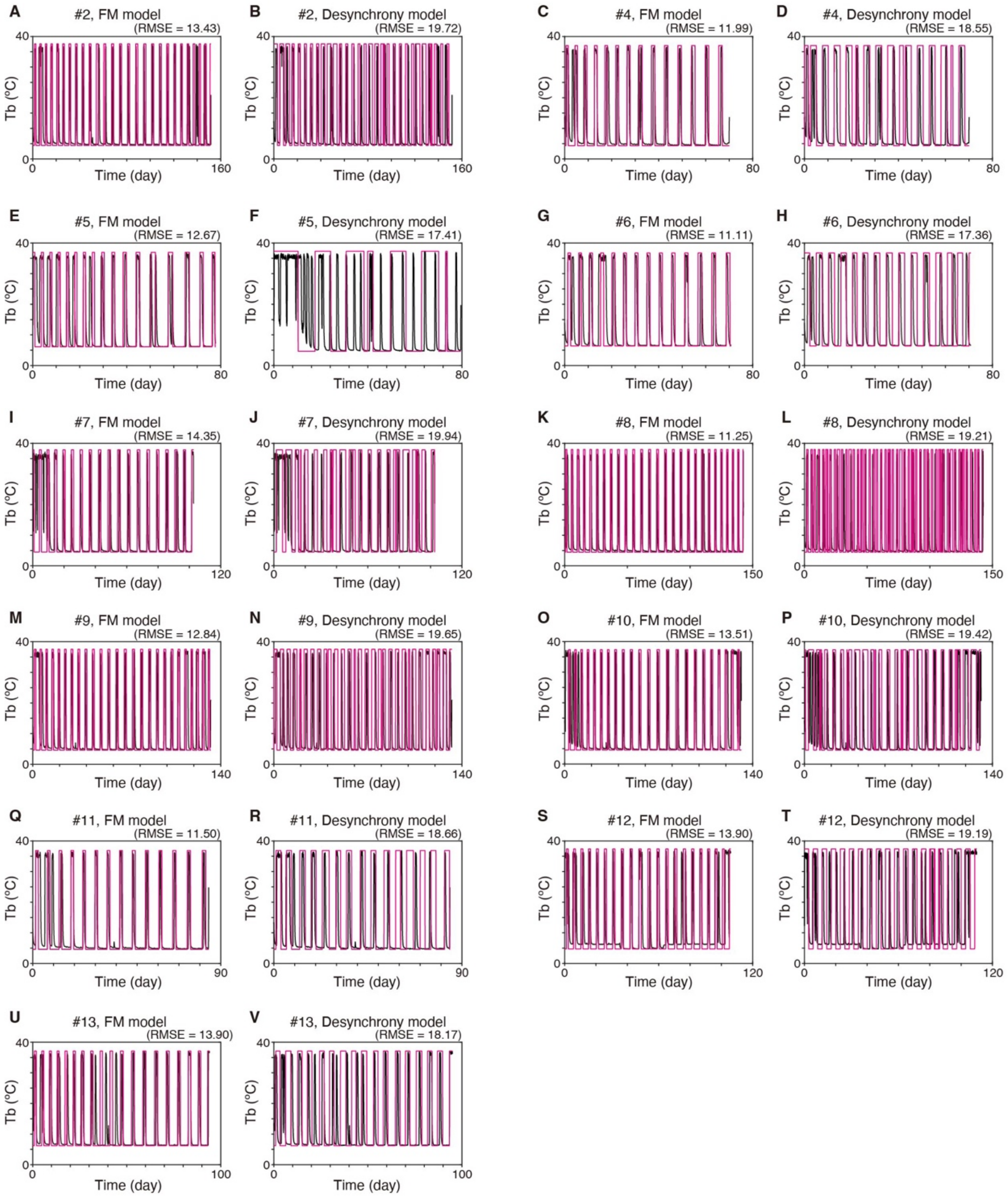


**Fig. S9. Frequency modulation and desynchrony model based time-evolution (red) best-fitted for recorded Tb data (black) of 11 animals using IS divergence (A-V).** The best-fitted parameter was chosen using IS divergence. Animal ID (#2, 4-13) is shown on the top of each graph. RMSE is the value of root mean squared error. Total 25 individual data including 2 in Fig. 2E, F, I, J and 12 in Fig. S10 were analyzed.

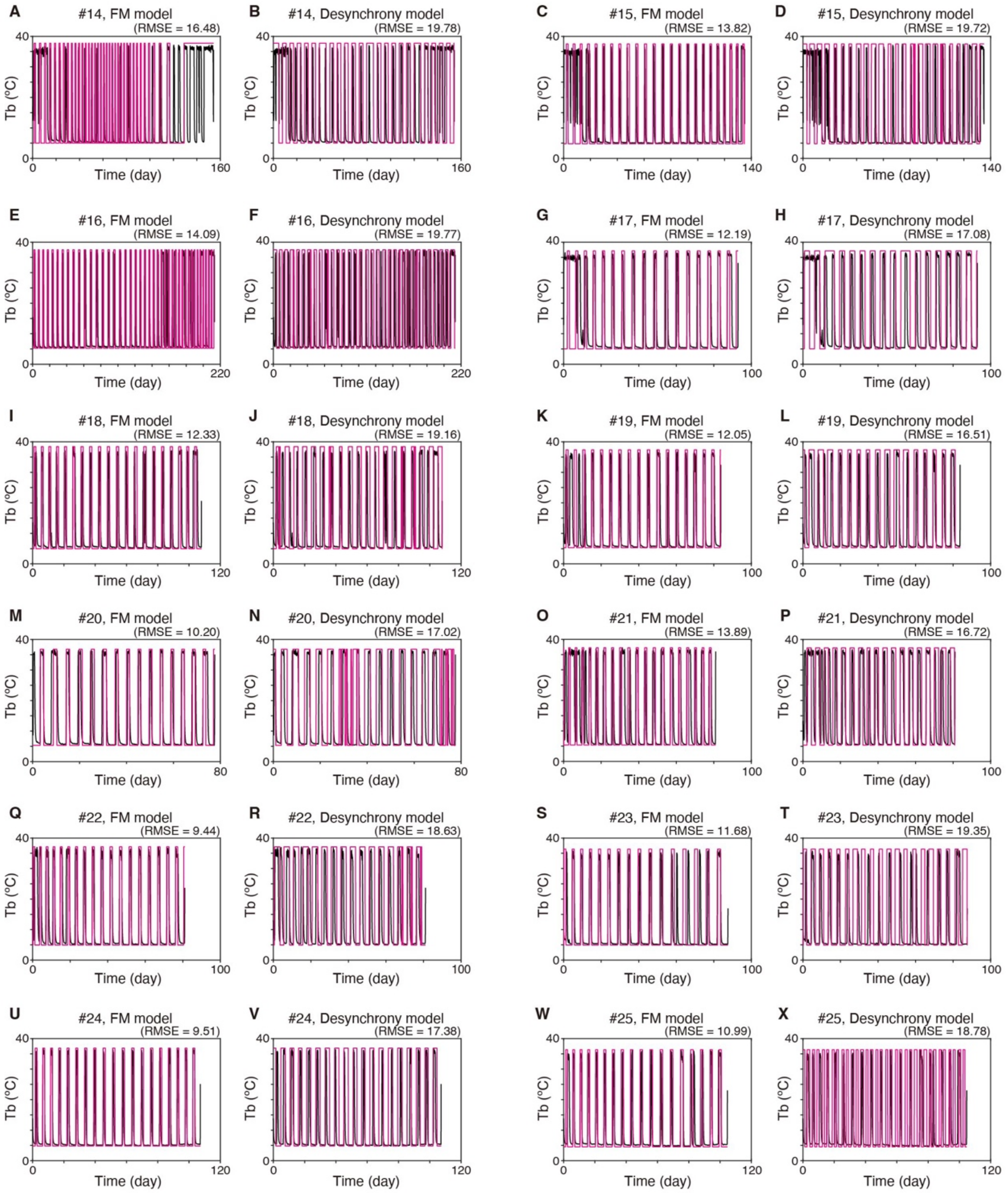


**Fig. S10. Frequency modulation and desynchrony model based time-evolution (red) best-fitted for recorded Tb data (black) of 12 animals using IS divergence (A-V).** The best-fitted parameter was chosen using IS divergence. Animal ID (#14-25) is shown on the top of each graph. RMSE is the value of root mean squared error. Total 25 individual data including 2 in Fig. 2E, F, I, J and 11 in Fig. S9 were analyzed.

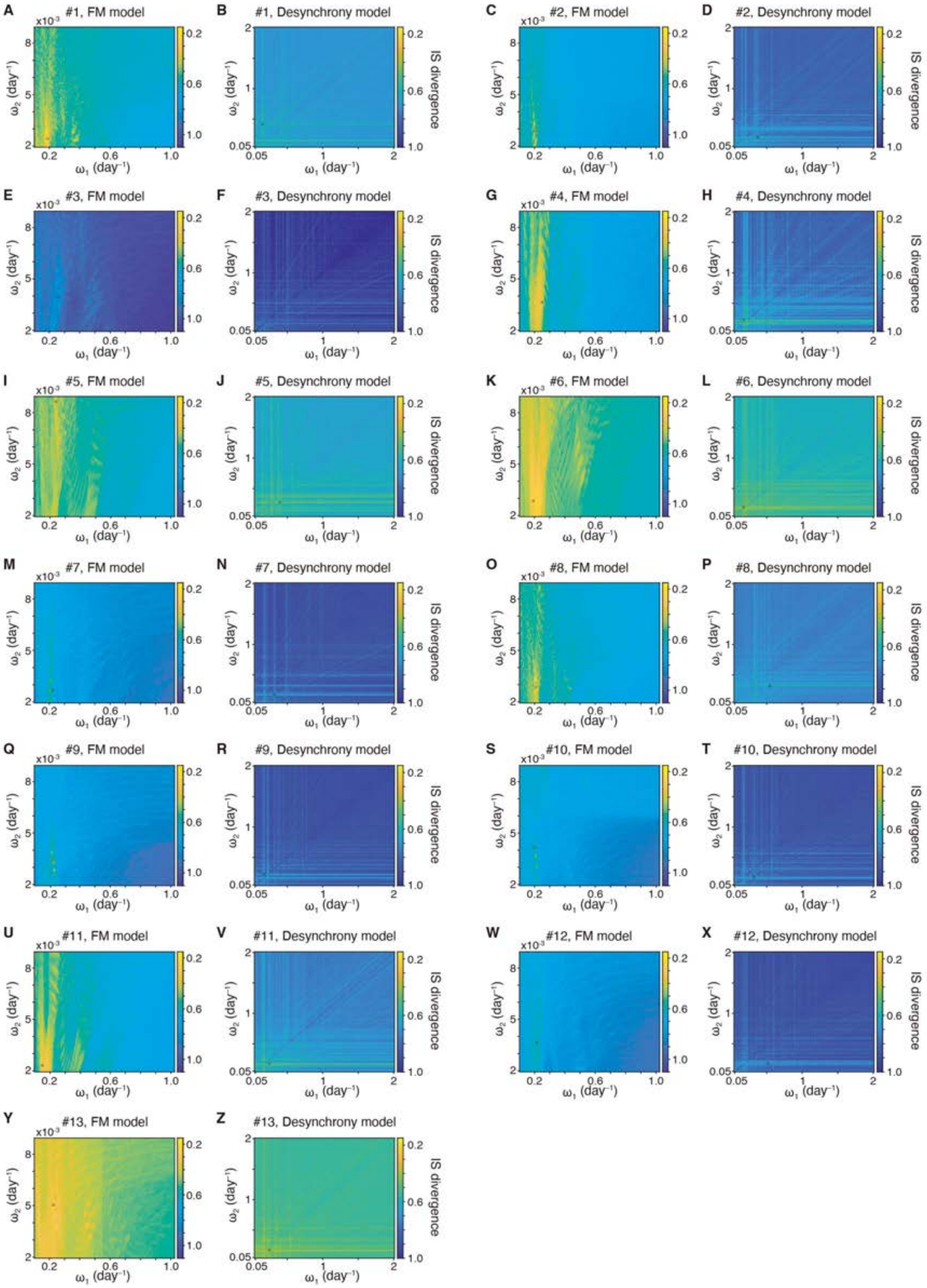


**Fig. S11. Distribution of IS divergence as a function of two frequencies ($\omega_1$ and $\omega_2$) of FM and desynchrony model for 13 individual experimental data (A-Z).** The best-fitted parameter, yielding the minimum IS divergence (circle) was used in Fig.2E, F, I, J, Fig. S9.

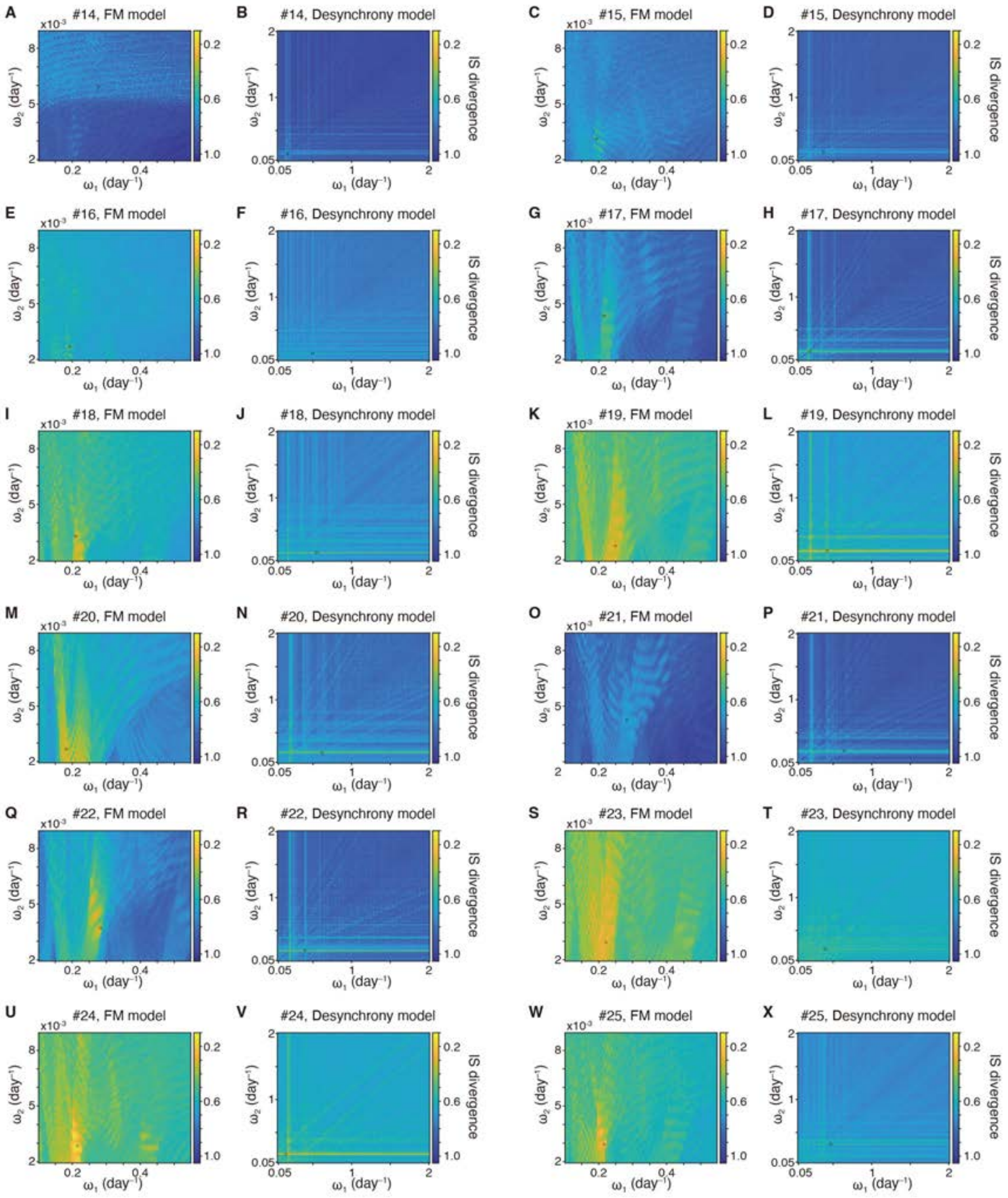


**Fig. S12. Distribution of IS divergence as a function of two frequencies ($\omega_1$ and $\omega_2$) of FM and desynchrony model for 12 individual experimental data (A-Z).** The best-fitted parameter, yielding the minimum IS divergence (circle) was used in Fig. S10.

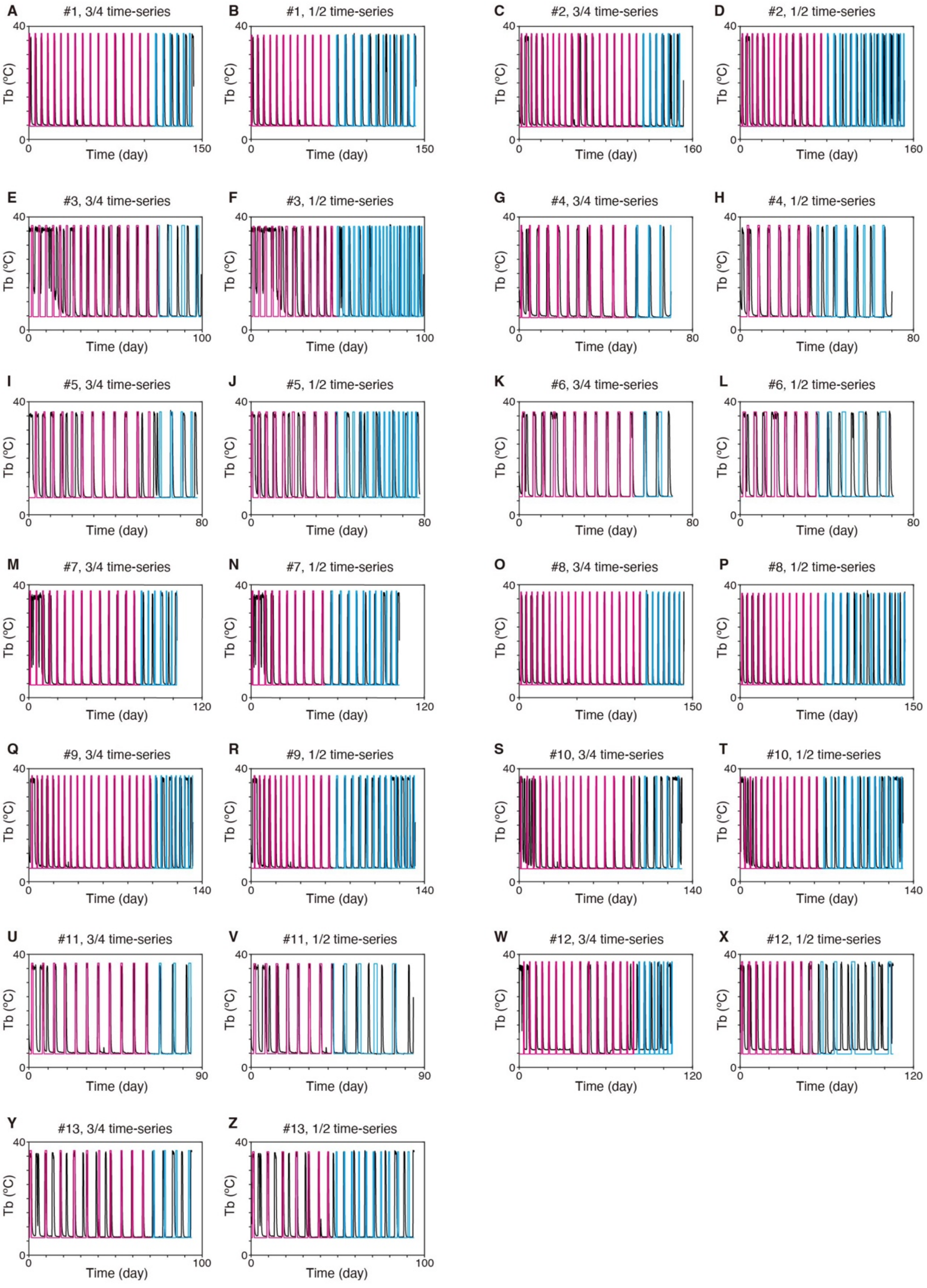


**Fig. S13. Forecasting Tb fluctuation from the three-quarters or the first half of the whole Tb time-series for 13 individual data using maximum likelihood (A-Z).** Tb time-series was reconstructed (magenta) and predicted (cyan). Recorded Tb data is shown by black.

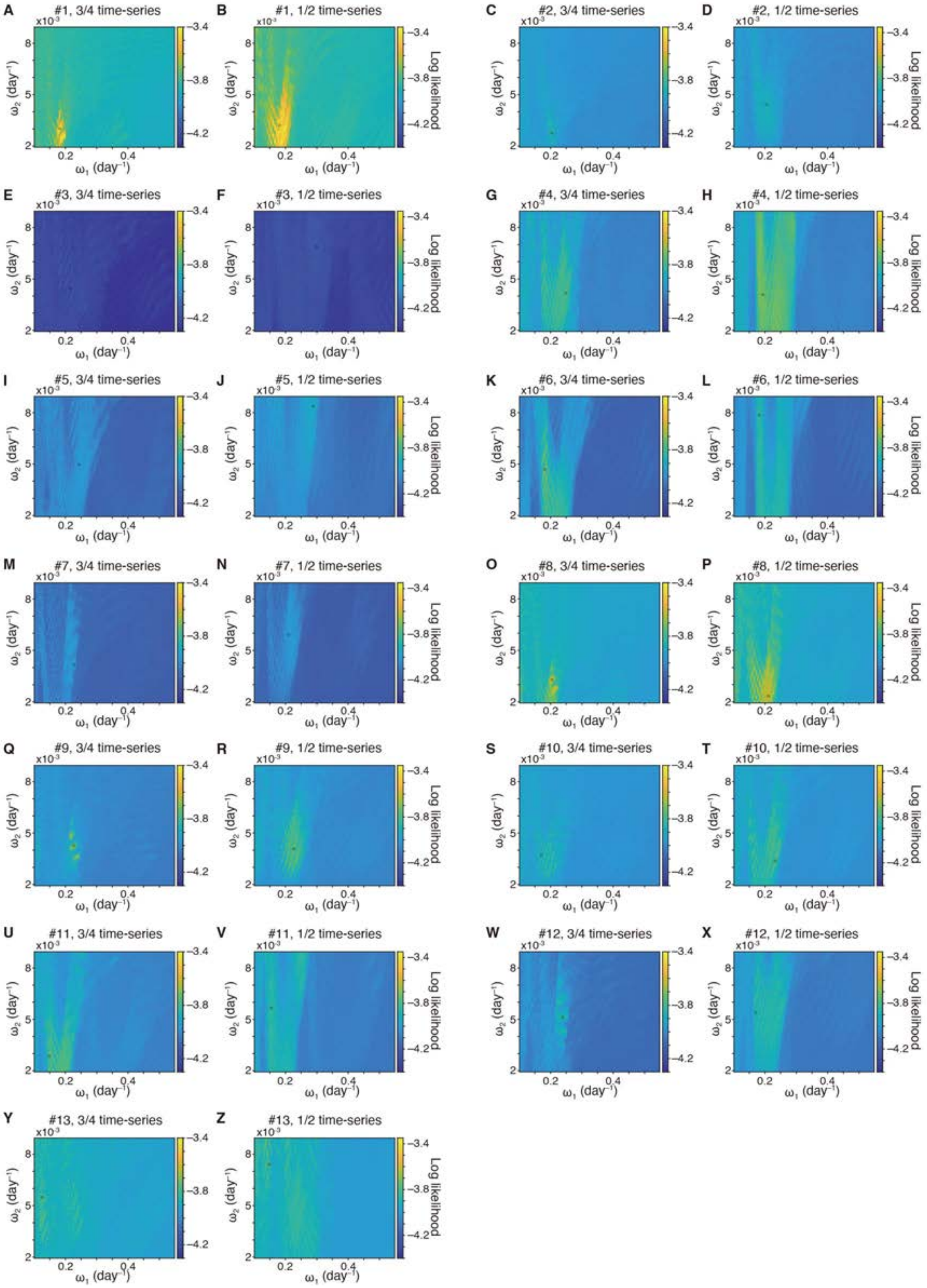

**Fig. S14. Distribution of log likelihood as a function of faster ($\omega_1$) and slower ($\omega_2$) frequency estimated from the three-quarters or the first half of the whole Tb time-series (A-Z).** The best-fitted parameter, yielding the maximum likelihood (circle) was used in Fig. S13.

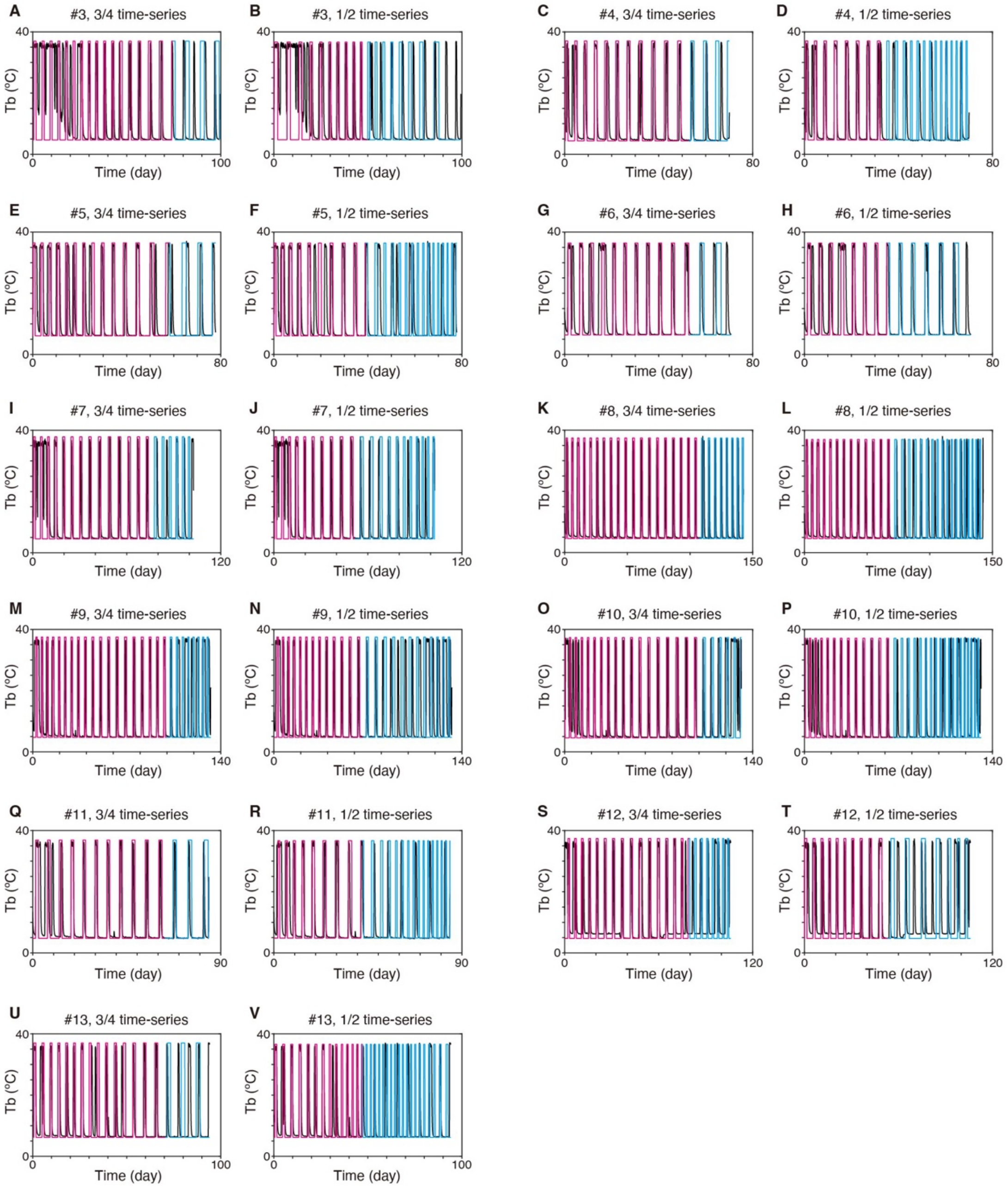


**Fig. S15. Forecasting Tb fluctuation from the three-quarters or the first half of the whole Tb time-series for 11 individual data using IS divergence (A-V).** Tb time-series was reconstructed (magenta) and predicted (cyan). Recorded Tb data is shown by black. Total 13 individual data including 2 in Fig. 4A-D were analyzed.

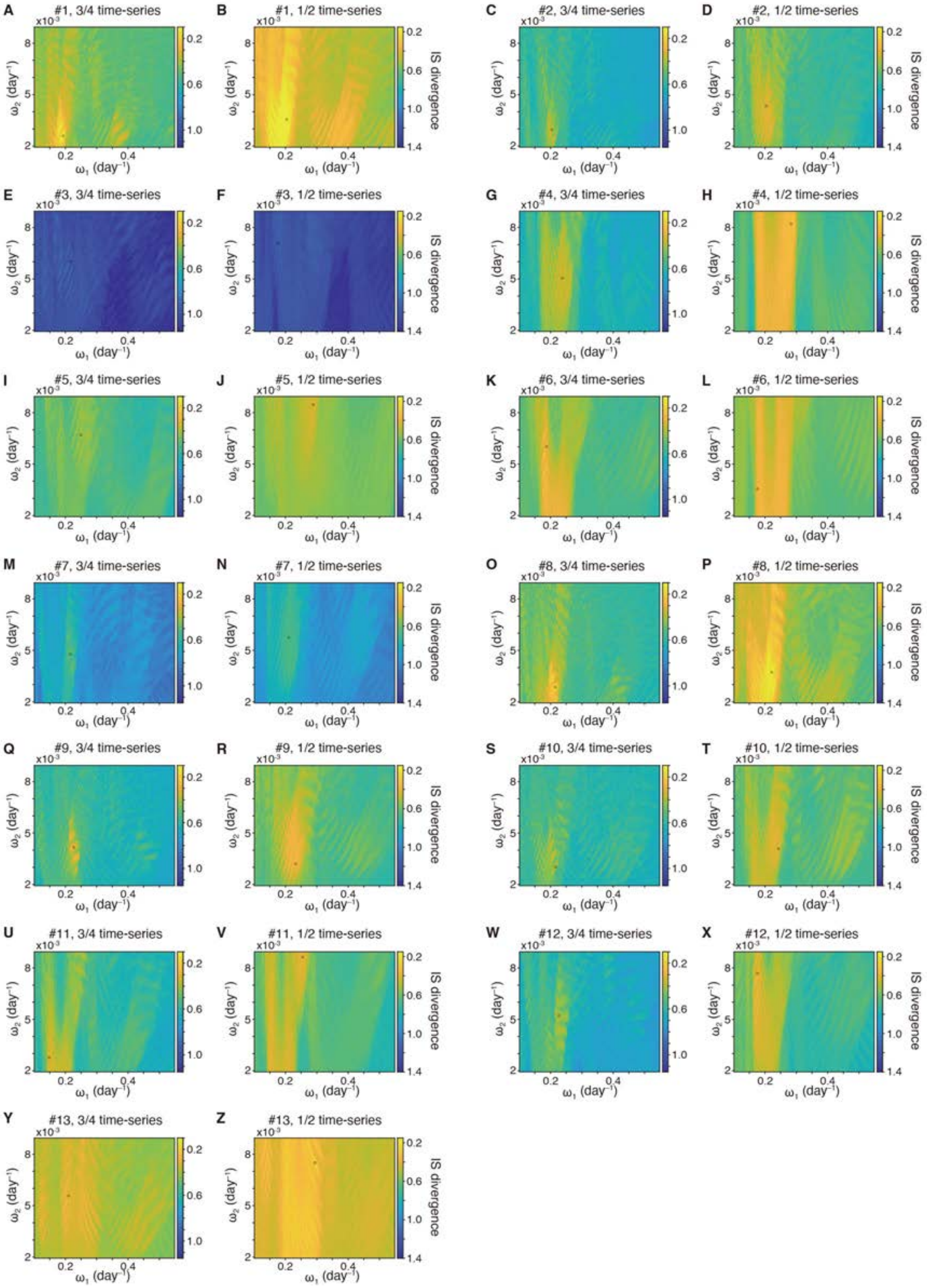


**Fig. S16. Distribution of IS divergence as a function of faster ($\omega_1$) and slower ($\omega_2$) frequency estimated from the three-quarters or the first half of the whole Tb time-series (A-Z).** The best-fitted parameter, yielding the minimum IS divergence (circle) was used in Fig. 4A-D, Fig. S15.

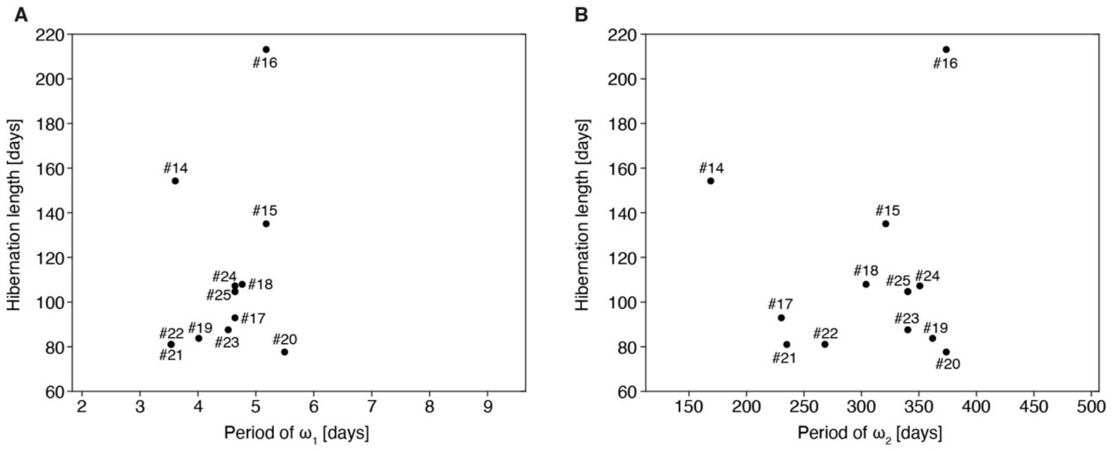


**Fig. S17. Distribution of hibernation length as a function of faster and slower period ($\omega_1$ and $\omega_2$ in FM model) for 12 individual experimental data (A,B).** The period of $\omega_1$ and $\omega_2$ were estimated using IS divergence. Total 12 individual data (#14-25) were quantified, for which Tb time-series of the whole hibernation period was measured. In our analysis, the offset of hibernation is defined to be the point from which Tb is higher than 15 degree for more than 10 days.

| Parameter | Lower limit | Upper limit | Number of intervals |
|---|---|---|---|
| $\omega_1$ | 0.104 /days | 1.02 /days | 165 |
| $\omega_2$ | 0.00197 /days | 0.00890 /days | 80 |
| $\phi_1$ | 0 | 2π | 20 |
| $\phi_2$ | 0 | 2π | 20 |
| $A_2$ | 0.124 | 0.206 | 20 |
| $\theta$ | 0.675 | 0.990 | 20 |

**Table S1. Parameter range for statistical analysis of frequency modulation model.** The upper and lower limits for parameter estimation were determined from preliminary estimation so that the model is likely to reproduce the torpor-IBA cycles of Syrian hamster.

| Parameter | Lower limit | Upper limit | Number of intervals |
|---|---|---|---|
| $\omega_1$ | 0.0500 /days | 2.00 /days | 80 |
| $\omega_2$ | 0.0500 /days | 2.00 /days | 80 |
| $\phi_1$ | 0 | 2π | 20 |
| $\phi_2$ | 0 | 2π | 20 |
| $A_2$ | 1 | 3 | 20 |
| $\theta$ | 0 | $\theta_{smax}$ | 20 |

**Table S2. Parameter range for statistical analysis of desynchrony model.** Parameter, $\theta$ is the threshold for step function *S* (see Eq. 2 for details). Desynchrony model can yield quasiperiodic oscillations with fluctuating amplitudes for certain parameter sets. The upper limit for the threshold, $\theta_{smax}$ was set to be the smallest local maximum, multiplied by 0.99 so that the small amplitudes of oscillations from desynchrony model were reflected in simulating Tb fluctuation.
